# A Compartmental Model for Simulating the Gut-Brain Axis in Gastric Function Regulation

**DOI:** 10.1101/2025.06.19.660643

**Authors:** Shannon Q Fernandes, Mayuresh V Kothare

**Affiliations:** Department of Chemical and Biomolecular Engineering, Lehigh University, Bethlehem, PA, 18015, USA

**Keywords:** gut-brain axis, vago-vagal loop, autonomic nervous system, computationally inexpensive model, compartmental modeling framework

## Abstract

Gastric function is regulated by the gut-brain axis, which integrates vagal and <u>enteric nervous system (ENS)</u> pathways. The parasympathetic circuit within the vagal pathway promotes digestion by stimulating peristaltic activity and relaxing the <u>Pyloric sphincter (PS)</u> through motor and sensory neurons. In contrast, the sympathetic pathway inhibits digestion by suppressing peristalsis and constricting the <u>PS,</u> highlighting the complex neural coordination involved in gastric regulation.

This study introduces a novel mathematical model of the gut-brain axis using a computationally efficient compartmental framework. The model simulates the vagal and <u>ENS</u> pathways and their corresponding effects on gastric function to improve our understanding of gut-brain axis regulation. The model employs the <u>Michaelis-Menten equation with a Hill coefficient (MMEHC)</u> equation to capture neurotransmitter release at neuromuscular junctions by stimulation of motor neurons and its effects on gastric cells. Motor (efferent) neurons are modeled for three key stomach regions: the fundus (tonic activity), antrum (phasic activity), and <u>PS</u> (both tonic and phasic activity). Thus, the stomach is represented as a three-compartment model.

The stomach model extends our previous work (Fernandes et al., 2024) by incorporating passive stress and dynamic changes in stomach geometry. Sensory (afferent) inputs are represented through linear equations that account for chemo- and mechanoreceptor activity, while a binary variable captures the sympathetic response. Afferent and efferent firing rates are linked via fitted curves to effectively close the gut-brain axis feedback loop, borrowing from a similar approach used to model cardiovascular regulation.

The simulation results align with physiological observations, demonstrating inhibitory digestive activity during sympathetic responses and excitatory activity, such as gastric emptying, during parasympathetic responses. During gastric emptying, the <u>Interstitial Cells of Cajal (ICC)</u> activity shows constant amplitude for low to medium gastric volumes but exhibits an increase in amplitude at very high gastric volumes. Furthermore, gastric emptying rates decrease with high-calorie liquids due to <u>PS</u> regulation, validating the potential of the model for studying <u>Gastrointestinal (GI)</u> disorders and developing vagal-based therapies.

## 1. Introduction and motivation

The neural control of the <u>GI</u> system integrates both extrinsic and intrinsic inputs. Gastric function is predominantly regulated by extrinsic inputs from the <u>central nervous system (CNS),</u> with the vagus nerve activity playing a pivotal role in mediating <u>CNS</u> influence on the stomach [1, 2, 3, 4, 5]. Intrinsic control mechanisms, including the submucosal and myenteric plexuses, as well as <u>ICC,</u> also contribute significantly to gastric motility regulation. The coordinated interplay among <u>ICC, Smooth Muscle Cells (SMC), ENS,</u> and vagal inputs establishes the patterns necessary for proper gastric function [6].

The <u>CNS</u> modulates gastric function via the dual influence of parasympathetic and sympathetic pathways, which cooperate to regulate digestive activities in the <u>GI.</u> These pathways primarily originate from neural circuits in the caudal brainstem. The sympathetic pathway involves cholinergic preganglionic neurons that originate in the intermediolateral column of the thoracic spinal cord. These neurons project to postganglionic neurons, which innervate the enteric plexus, a localized neural network within the stomach. The sympathetic pathway primarily modulates gastric inhibition by suppressing cholinergic vagal inputs to postganglionic neurons [2, 7, 8]. This pathway is activated during “fight or flight” responses, highlighting its role in stress-induced gastric modulation [9].

The parasympathetic pathway employs both excitatory and inhibitory signaling to regulate various gastric functions, including promoting gastric emptying [10]. Parasympathetic control is mediated through the <u>dorsal motor nucleus of the vagus (DMV),</u> which governs the vagovagal reflex. The parasympathetic neurons in the <u>DMV</u> consist of cholinergic preganglionic neurons projecting to postganglionic neurons within the enteric plexus. These postganglionic neurons, which can be either cholinergic or <u>non-adrenergic, non-cholinergic (NANC),</u> provide excitatory or inhibitory signals to the stomach.

Sensory signaling from the gut to the brainstem involves mechano- and chemo-sensitive inputs transmitted via vagal afferent fibers to the <u>nucleus tractus solitarius (NTS).</u> The <u>NTS</u> processes these inputs and relays them to the <u>DMV</u> through direct or indirect pathways. Indirect pathways integrate signals from limbic and hypothalamic regions, modulating the reflexive output to regulate gastric function [11, 12]. Vagal efferent fibers from the <u>DMV</u> project back to the gut, controlling gastric motility. This bidirectional communication between the gut and brainstem, known as the vagovagal reflex, is crucial for maintaining gastric homeostasis [6].

Notably, the synaptic connections between the <u>NTS</u> and <u>DMV</u> in the brainstem are not static but exhibit plasticity. Previous studies [10, 13, 14, 15, 2] have shown that this plasticity allows the synaptic circuitry to adapt and fine-tune gastric motor activity in both physiological and pathological states, depending on sensory input. This adaptability underscores the complexity and significance of neural control in gastric function.

The vagovagal reflex is crucial, as disruptions in vagal sensory-motor function can lead to <u>GI</u> disorders. Evidence of this was reported in [16, 17, 18], where patients with functional dyspepsia exhibited altered vagal function characterized by reduced gastric compliance and impaired gastric emptying. These findings suggest that impaired vagal regulation may underlie many functional gastrointestinal disorders [2]. Despite growing recognition of the gut-brain axis and its influence on gastric control, current mathematical models in the literature lack a comprehensive representation of the vagal pathways involved and their regulatory influence on gastric function.+

This study aims to develop a computationally efficient mathematical model of the gut-brain axis neural circuitry governing the gastric function. This model will integrate seamlessly with our previously established compartmental framework for gastric function [19] and provide insights into the dynamic interactions between neural and gastric systems. Moreover, such a model holds great potential for its use in advancing therapeutic interventions such as vagal nerve stimulation for treating gastric diseases using concepts from model-based control theory [20].

## 2. Methods: Formulating the mathematical model

### 2.1. Modeling the autonomic nervous system (ANS) regions involved in gastric function regulation: An overview

The mathematical model of the vagal brain-gut axis circuitry controlling gastric function is structured into several components, which are discussed in the following subsections. The first subsection outlines the efferent vagal pathways, which mediate gastric motor responses through excitatory and inhibitory signaling. The second subsection derives equations for afferent pathways, capturing sensory input from chemo-and mechanoreceptors. The third subsection examines the interneuronal connections within the brain, specifically between the afferent and efferent vagal fibers, for instance, the <u>NTS</u> and <u>DMV</u> interneuron connection. Finally, the fourth subsection integrates the derived pathways to model the parasympathetic and sympathetic circuits governing gastric motility and emptying, offering a unified framework for the gut–brain axis.

The stomach is modeled using a compartmental framework as proposed in our previous study [19]. In this study, the framework divides the stomach into three compartments: the fundus, antrum, and <u>PS,</u> represented by the subscripts *w* = 1, 2, 3, respectively. Unlike the fixed geometry assumed in the prior model, this study introduces dynamic compartmental geometry, which accounts for changes in gastric volume during motility and emptying. This enhancement provides a more realistic representation of gastric behavior.

Passive stress models, dependent on gastric volume, are integrated into the compartmental framework to capture the biomechanical properties of the stomach. The dynamic volume is constrained between 0.08 L (empty stomach) and 1.2 L (full stomach), consistent with physiological observations reported in prior studies [21, 22, 23, 24, 25, 26].The parameters for the model used in this study are summarized in Tables 1 and 2. For ease of readability, the main equations are included in the paper and the mathematical details of the modeling steps are provided in the supplementary material section which is referred to as Section 8.

**Table 1.**
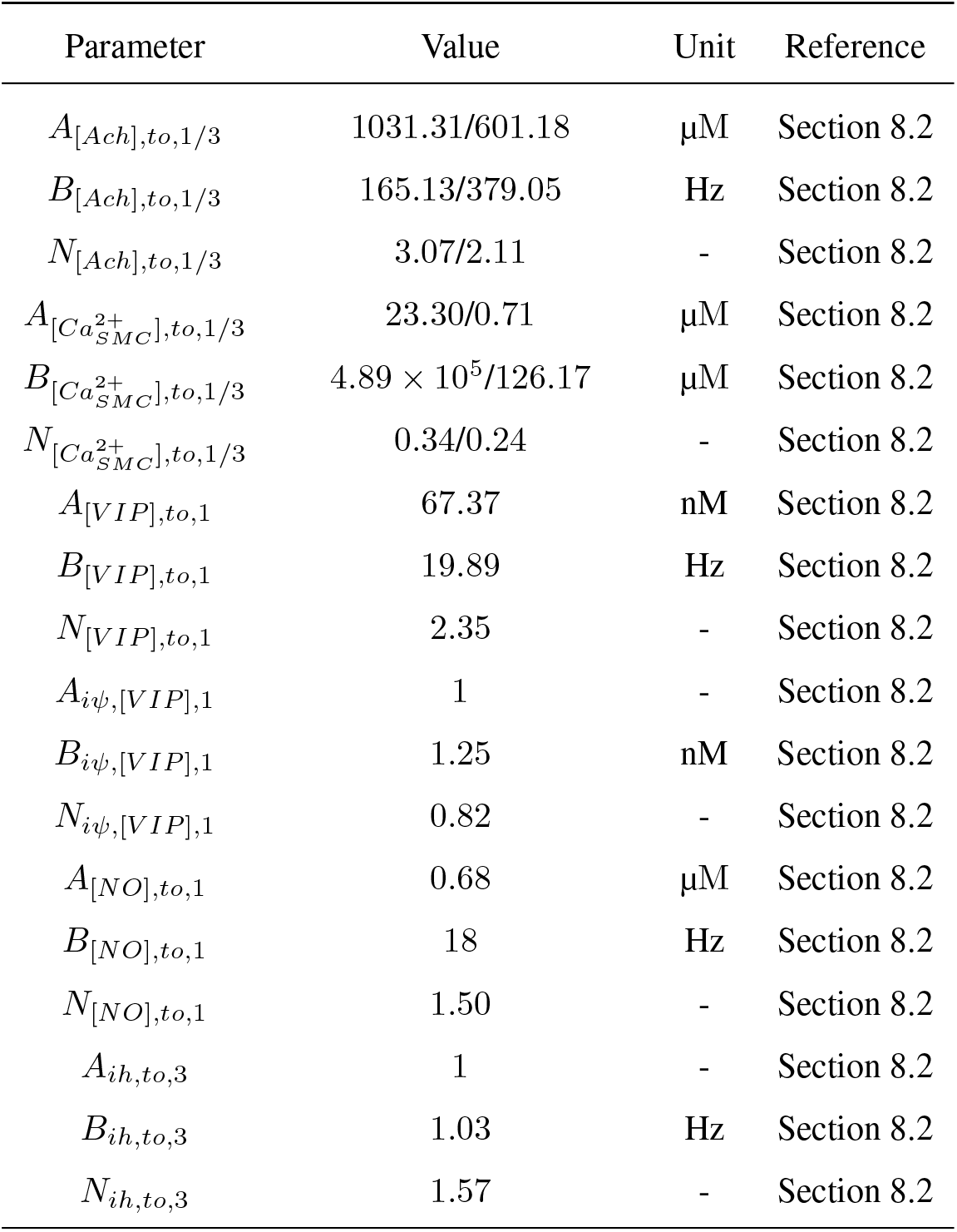
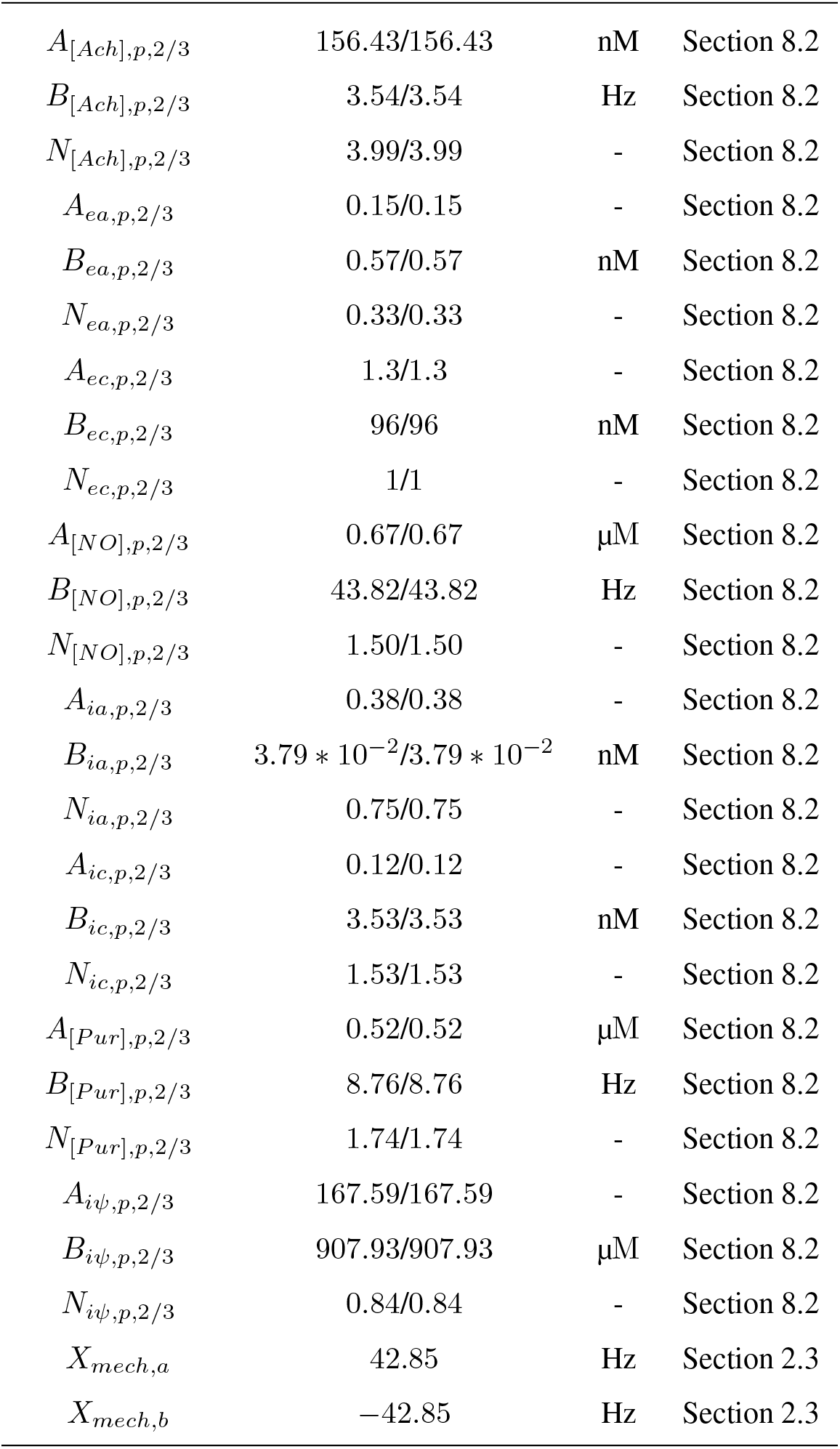
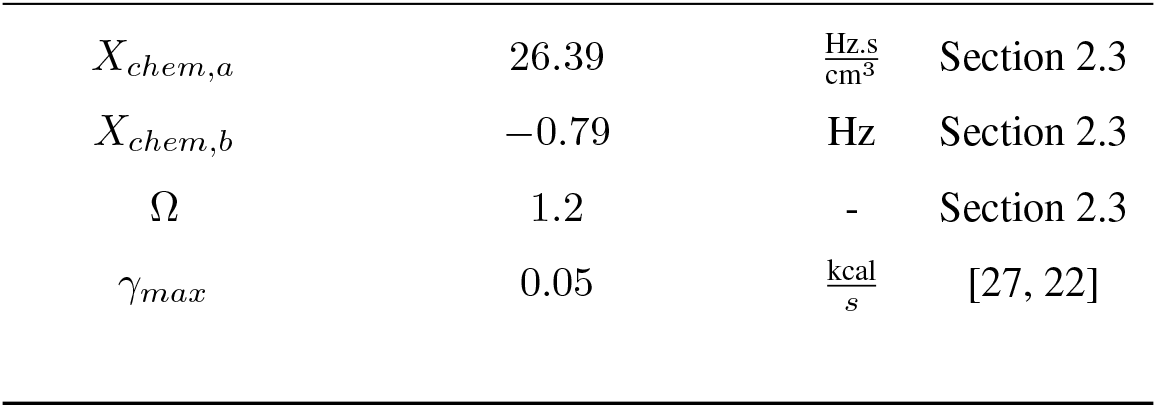
Neural model parameters for simulating gut-brain axis for gastric function regulation.

| Parameter | Value | Unit | Reference |
| --- | --- | --- | --- |
| $A_{[Ach],to,1/3}$ | 1031.31/601.18 | $\mu\text{M}$ | Section 8.2 |
| $B_{[Ach],to,1/3}$ | 165.13/379.05 | Hz | Section 8.2 |
| $N_{[Ach],to,1/3}$ | 3.07/2.11 | - | Section 8.2 |
| $A_{[Ca_{SMC}^{2+}],to,1/3}$ | 23.30/0.71 | $\mu\text{M}$ | Section 8.2 |
| $B_{[Ca_{SMC}^{2+}],to,1/3}$ | $4.89 \times 10^5/126.17$ | $\mu\text{M}$ | Section 8.2 |
| $N_{[Ca_{SMC}^{2+}],to,1/3}$ | 0.34/0.24 | - | Section 8.2 |
| $A_{[VIP],to,1}$ | 67.37 | nM | Section 8.2 |
| $B_{[VIP],to,1}$ | 19.89 | Hz | Section 8.2 |
| $N_{[VIP],to,1}$ | 2.35 | - | Section 8.2 |
| $A_{i\psi,[VIP],1}$ | 1 | - | Section 8.2 |
| $B_{i\psi,[VIP],1}$ | 1.25 | nM | Section 8.2 |
| $N_{i\psi,[VIP],1}$ | 0.82 | - | Section 8.2 |
| $A_{[NO],to,1}$ | 0.68 | $\mu\text{M}$ | Section 8.2 |
| $B_{[NO],to,1}$ | 18 | Hz | Section 8.2 |
| $N_{[NO],to,1}$ | 1.50 | - | Section 8.2 |
| $A_{ih,to,3}$ | 1 | - | Section 8.2 |
| $B_{ih,to,3}$ | 1.03 | Hz | Section 8.2 |
| $N_{ih,to,3}$ | 1.57 | - | Section 8.2 |
| *Continued on next page* |  |  |  |
| $A_{[Ach],p,2/3}$ | 156.43/156.43 | nM | Section 8.2 |
| $B_{[Ach],p,2/3}$ | 3.54/3.54 | Hz | Section 8.2 |
| $N_{[Ach],p,2/3}$ | 3.99/3.99 | - | Section 8.2 |
| $A_{ea,p,2/3}$ | 0.15/0.15 | - | Section 8.2 |
| $B_{ea,p,2/3}$ | 0.57/0.57 | nM | Section 8.2 |
| $N_{ea,p,2/3}$ | 0.33/0.33 | - | Section 8.2 |
| $A_{ec,p,2/3}$ | 1.3/1.3 | - | Section 8.2 |
| $B_{ec,p,2/3}$ | 96/96 | nM | Section 8.2 |
| $N_{ec,p,2/3}$ | 1/1 | - | Section 8.2 |
| $A_{[NO],p,2/3}$ | 0.67/0.67 | μM | Section 8.2 |
| $B_{[NO],p,2/3}$ | 43.82/43.82 | Hz | Section 8.2 |
| $N_{[NO],p,2/3}$ | 1.50/1.50 | - | Section 8.2 |
| $A_{ia,p,2/3}$ | 0.38/0.38 | - | Section 8.2 |
| $B_{ia,p,2/3}$ | $3.79 * 10^{-2}/3.79 * 10^{-2}$ | nM | Section 8.2 |
| $N_{ia,p,2/3}$ | 0.75/0.75 | - | Section 8.2 |
| $A_{ic,p,2/3}$ | 0.12/0.12 | - | Section 8.2 |
| $B_{ic,p,2/3}$ | 3.53/3.53 | nM | Section 8.2 |
| $N_{ic,p,2/3}$ | 1.53/1.53 | - | Section 8.2 |
| $A_{[Pur],p,2/3}$ | 0.52/0.52 | μM | Section 8.2 |
| $B_{[Pur],p,2/3}$ | 8.76/8.76 | Hz | Section 8.2 |
| $N_{[Pur],p,2/3}$ | 1.74/1.74 | - | Section 8.2 |
| $A_{i\psi,p,2/3}$ | 167.59/167.59 | - | Section 8.2 |
| $B_{i\psi,p,2/3}$ | 907.93/907.93 | μM | Section 8.2 |
| $N_{i\psi,p,2/3}$ | 0.84/0.84 | - | Section 8.2 |
| $X_{mech,a}$ | 42.85 | Hz | Section 2.3 |
| $X_{mech,b}$ | -42.85 | Hz | Section 2.3 |
| $X_{chem,a}$ | 26.39 | $\frac{\text{Hz.s}}{\text{cm}^3}$ | Section 2.3 |
| $X_{chem,b}$ | -0.79 | Hz | Section 2.3 |
| $\Omega$ | 1.2 | - | Section 2.3 |
| $\gamma_{max}$ | 0.05 | $\frac{\text{kcal}}{\text{s}}$ | [27, 22] |

**Table 2.**
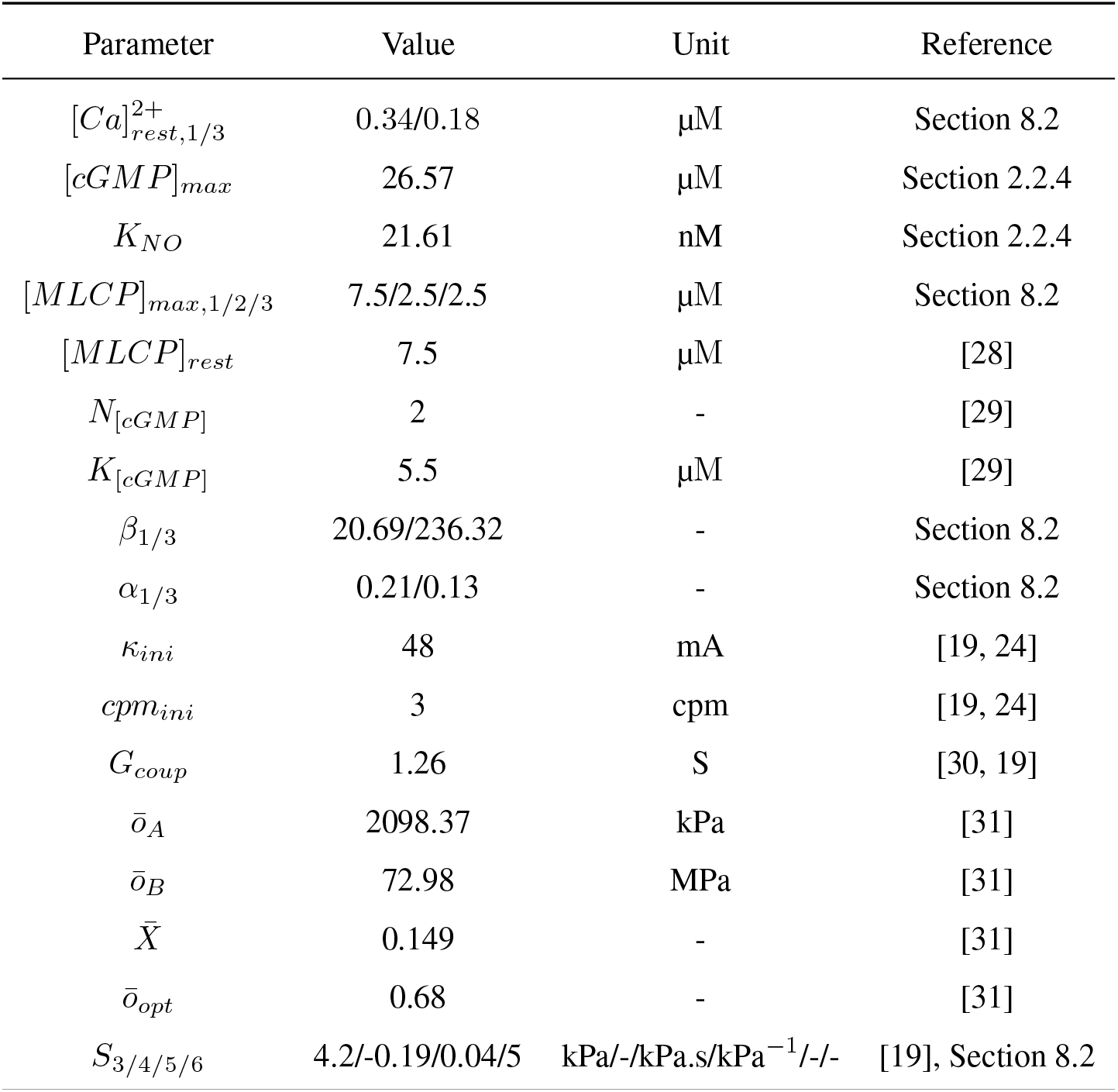
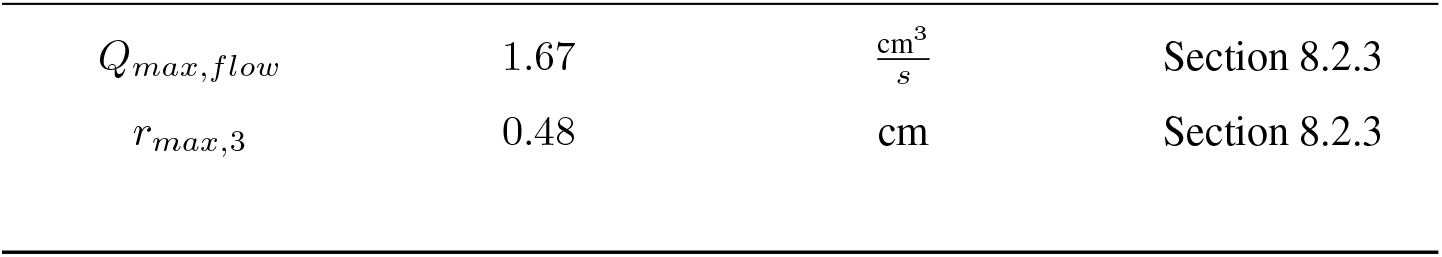
Gastric model parameters for simulating gut-brain axis for gastric function regulation.

### 2.2. Modeling the efferent/motor neuron connections and the stomach compartments

From the literature [32, 33, 34, 35, 36, 37, 38, 39, 23, 19, 22], gastric function varies significantly between three primary regions of the stomach: the fundus, antrum, and <u>PS.</u> Consequently, a three-compartment model is considered for this study.

The fundus (proximal stomach) primarily facilitates storage and gastric accommodation through distention [32, 33]. The antrum exhibits peristaltic activity, which drives fluid motion within the stomach, playing a crucial role in gastric mixing and emptying [21, 19, 23]. The <u>PS</u> functions as a valve, regulating the flow of gastric contents to the duodenum based on factors such as the caloric content of the meal [22, 40].

This understanding allows for the differentiation of motor neuron targets specific to each stomach region. Additionally, contraction patterns vary across these regions: the fundus primarily exhibits tonic activity, the antrum displays phasic activity, and the <u>PS</u> demonstrates both tonic and phasic activity [41, 42, 43, 44, 45, 24].

In the subsequent subsections, motor neuron activity—both inhibitory and excitatory—will be discussed, and mathematical formulations for these activities will be derived for each of these stomach regions.

#### 2.2.1. Motor neurons influencing tonic activity: Fundus

Unlike other regions of the stomach, the fundus (or proximal stomach) lacks <u>ICC,</u> which are essential to generate phasic contractions. Consequently, the fundus primarily maintains basal tone [24, 41]. The <u>NANC</u> inhibitory pathway reduces basal tone and serves as a critical mechanism for fundic relaxation [46]. In contrast, the cholinergic excitatory pathway increases basal tone and facilitates fundic contractions [47].

The <u>NANC</u> inhibitory pathway mediates fundic relaxation through the release of <u>Nitric oxide (NO)</u> and <u>Vasoactive intestinal peptide (VIP)</u> as primary neurotransmitters. At low neuronal firing frequencies, <u>NO</u> is released at the neuromuscular junction to induce relaxation [48, 49, 50]. At higher neuronal firing frequencies, <u>NO</u> and <u>VIP</u> are released simultaneously, further enhancing the relaxation response [51].

The cholinergic excitatory pathway induces the release of <u>Acetylcholine (Ach)</u> at the neuromuscular junction, promoting an increase in fundic contractions [52]. A schematic representation of the excitatory and inhibitory pathways in the fundus (*w* = 1) is illustrated in Fig. 1.

**Figure 1.**
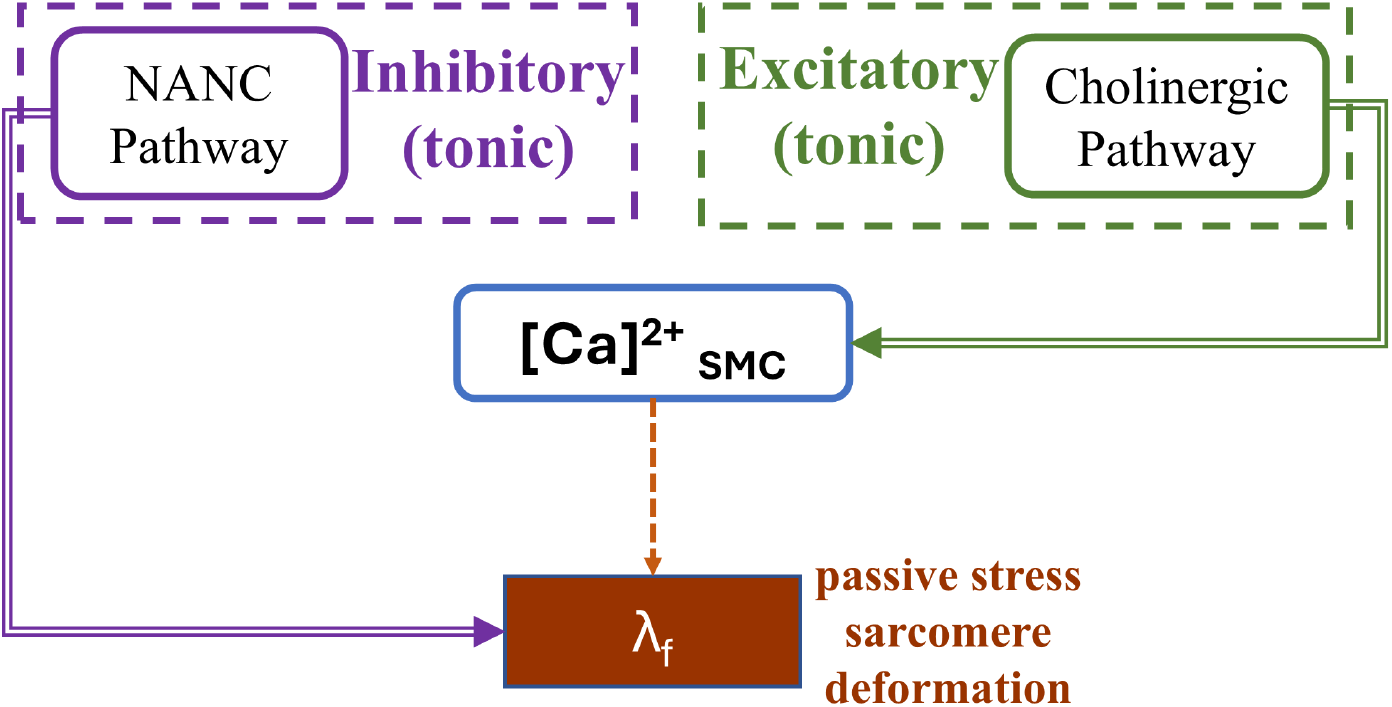
Schematic representation of excitatory and inhibitory inputs to for fundic tonic activity

To model the relationship between motor neuron firing frequency and neurotransmitter release at the neuromuscular junction, a <u>MMEHC</u> is employed. This equation effectively models the trends reported in a previous study on neuron firing and neurotransmitter release at the neuromuscular junction [53].

The inclusion of the Hill coefficient allows the equation to account for the nonlinear behavior often observed in neurotransmitter release dynamics. Additionally, the Michaelis-Menten formulation incorporates a saturation term, which is critical for representing the physiological limit of neurotransmitter release.

For simulating neurotransmitter receptor signaling in smooth muscle, the <u>MMEHC</u> equation is equally suitable. It is widely used to model lumped ligand-receptor interactions and is therefore employed to describe both neurotransmitter release and subsequent receptor-mediated signaling behaviors [54, 55]. Further details on the application of the <u>MMEHC</u> equation is discussed in Section 8.1.

The general form of the <u>MMEHC</u> equation incorporating a Hill coefficient is given by

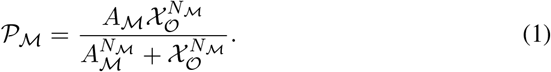

In this equation, *A* denotes the maximum response, *B* is the Michaelis constant, and *N* is the Hill coefficient. The subscript ℳ refers to the modality of the interaction, such as neurotransmitter type, inhibitory or excitatory activity, tonic (*to*) or phasic (*p*) contraction type, or compartment index *w*. The variable 𝒳 represents the input signal, which may correspond to a neurotransmitter concentration, a signaling molecule concentration, or the firing frequency of excitatory or inhibitory neurons. The subscript *O* designates the origin or context of this input, including notations such as *e* (excitatory), *i* (inhibitory), *to* (tonic), *p* (phasic), or compartment index *w*.

In this study, multiple equations—specifically Eqs. 2 to 6, 8, 9, 17 to 24 and 44 are formulated based on the generalized <u>MMEHC</u> expression and follow a similar form.

#### 2.2.2. Cholinergic pathway (Ach): Fundus

In the fundus, the neurotransmitter <u>Ach</u> is released at the neuromuscular junction via the cholinergic pathway [47]. Acetylcholine increases intracellular calcium concentration 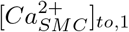 in the smooth muscle, influencing fundic contractions [53]. Consequently, the <u>MMEHC</u> equation is formulated to describe the concentration of *Ach* released, which depends on the firing frequency of the cholinergic pathway *f*_*e*,*to*,1_. This relationship is expressed as follows

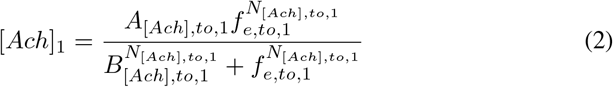

The <u>MMEHC</u> equation for the intracellular calcium concentration in the <u>SMC</u> 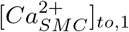 influenced by tonic cholinergic neurotransmitter signaling is given as

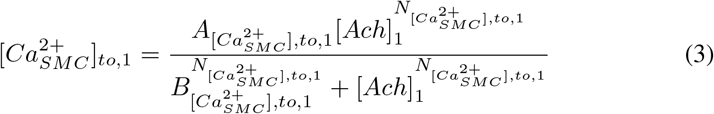

Data from Kim et al., 2020 and da Silva et al., 2018 [52, 56] were used to fit the parameters of the cholinergic signaling response for Eqs. <u>2</u> and <u>3.</u>

#### 2.2.3. NANC pathway (NO and VIP): Fundus

The neurotransmitters <u>NO</u> and <u>VIP</u> are released at the neuromuscular junction of the fundus via <u>NANC</u> pathway signaling [57]. Based on the firing frequency of the <u>NANC</u> pathway *f*_*i*,*to*,1_, the concentration of the neurotransmitter <u>VIP</u> released at the neuromuscular junction is described by the <u>MMEHC</u> equation as follows

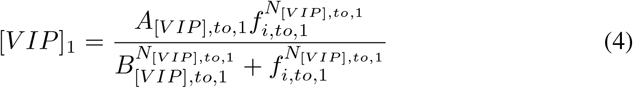

The <u>VIP</u> neurotransmitter is known to inhibit <u>myosin light chain kinase (MLCK)</u> concentration via the <u>cyclic adenosine monophosphate (cAMP)</u> signaling pathway [29, 58]. The <u>MLCK</u> concentration inhibitory factor *ϕ*_*iψ*,1_ is modeled using the <u>MMEHC</u> equation and is expressed as follows

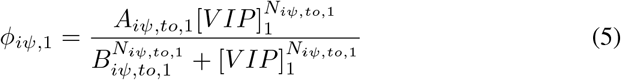

Based on the firing frequency *f*_*i*,*to*,1_, the release of the neurotransmitter <u>NO</u> at the neuromuscular junction is represented by the <u>MMEHC</u> equation as follows

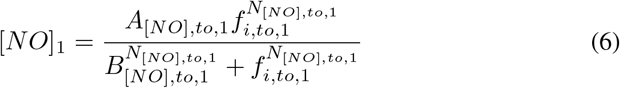

Data from previous studies [59, 60, 61, 46] were used to fit the <u>NO-mediated</u> neurotransmitter inhibitory response through increased <u>myosin light chain phosphatase (MLCP)</u> concentration, which will be discussed in the following section. Additionally, data from [59, 62, 46, 60, 63] were used to fit the inhibitory response associated with the reduction in <u>MLCK</u> concentration.

#### 2.2.4. Equations in the compartmental model for Fundus

The total intracellular calcium concentration in the fundus <u>SMC</u> 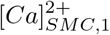 is the sum of the resting calcium concentration 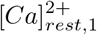 in the fundus and the increase in calcium concentration influenced by cholinergic signaling 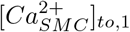. The relationship is expressed as

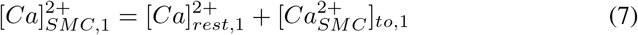

The Gajendiran and Buist, 2010 model [28] is employed to describe the kinetics that convert intracellular calcium concentration into active <u>MLCK</u> [*MLCK*_*act*_]_*to*,1_.

<u>NO</u> signaling increases <u>MLCP</u> concentration via the <u>cyclic guanosine monophosphate (cGMP)</u> pathway, as demonstrated in studies [64, 29]. Data from Yang et al., 2005 [29] were used to fit a Michaelis-Menten kinetic equation modeling the relationship between <u>NO</u> and <u>cGMP.</u> The <u>cGMP</u> concentration [*cGMP*]_*to*,1_ in the fundus is represented as

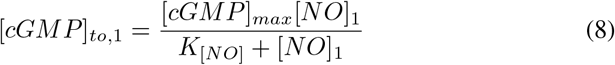

The <u>MMEHC</u> model presented in Yang et al., 2005 [29] was applied to model the relationship between <u>cGMP</u> and <u>MLCP.</u> The increase in <u>MLCP</u> concentration [*MLCP*]_*f*,*to*,1_ in the fundus is described by the following equation

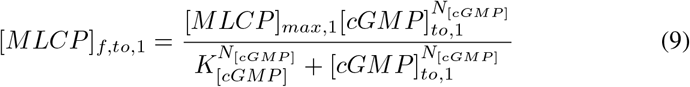

The total <u>MLCP</u> concentration is given as [*MLCP*]_*to*,1_ = [*MLCP*]_*rest*_+[*MLCP*]_*f*,*to*,1_. Similarly, the total activated <u>MLCK</u> concentration is expressed as [*MLCK*]_*to*,1_ = *ϕ*_*iψ*,1_[*MLCK*_*act*_]_*to*,1_ where *ϕ*_*iψ*,1_ represents the inhibitory factor induced by <u>VIP</u> neurotransmitter signaling, affecting <u>MLCK</u> concentration.

The Hai-Murphy model [65], as reported in the Gajendiran and Buist, 2011 paper [28], is utilized to compute the total number of latch bridges formed. Tonic contractions, being sustained contractions, result in the formation of latch bridges as crossbridge states which transition into latch bridges during sustained contractions [66].

The total number of latch bridges is represented as the sum of [*AM*_*p*_]_*to*,1_ and [*AM*]_*to*,1_. The equations for modeling the formation of latch bridges based on the Hai-Murphy model [65] are given as follows

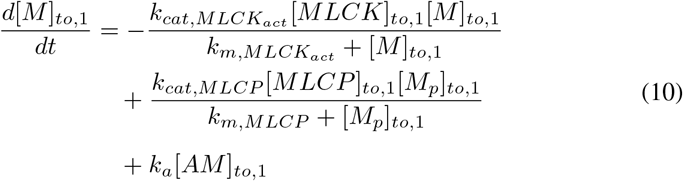

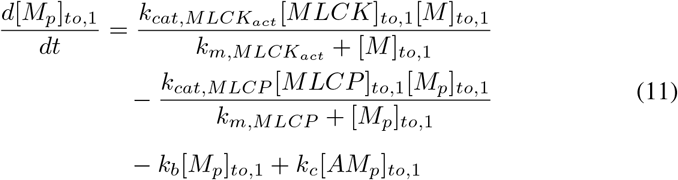

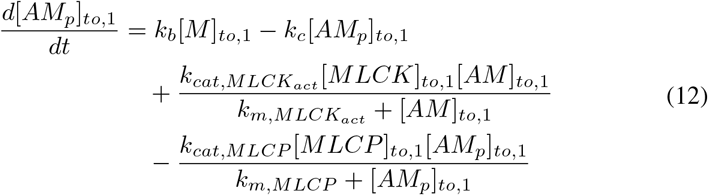

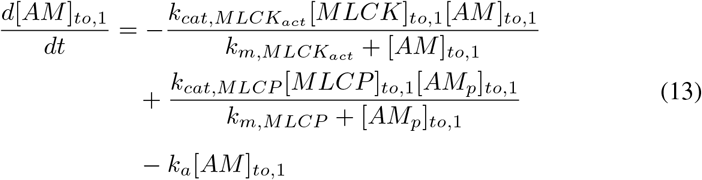

The relative area for the distension of the stomach in the fundus compartment *RA*_1_ as a function of the total number of latch bridges is calculated using the equation reported in Wang et al, 2008 [67], expressed as

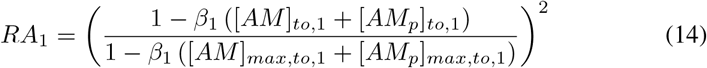

Here *β*_1_ is a dimensionless parameter. Considering the stomach can be approximated as a cylinder [19], the radius *r*_*fin*_ of the fundus compartment is computed using the open cylinder equation as

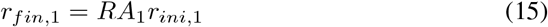

The radius of the open cylinder fundus at its minimum volume, *r*_*ini*,1_, corresponds to the unstressed radius of the stomach, calculated when the gastric volume is at a minimum of 0.08 L.

The sarcomere deformation of the muscle fiber *λ*_*f*,1_, which influences the passive stress component in the fundus wall, is determined by a linear relationship involving the ratio of the stressed radius to the unstressed radius, as defined in the study by Pironet et al., 2013 [68]. The dimensionless constant *α*_1_ is introduced to fit *λ*_*f*,1_ within the physiological range.

The equation for the passive stress sarcomere deformation *λ*_*f*,1_ is expressed as

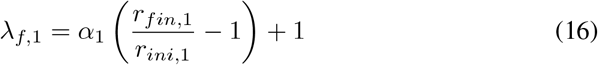

#### 2.2.5. Motor neurons influencing phasic activity: Antrum

The antrum of the stomach plays a pivotal role in mixing and grinding ingested food [19, 21]. It is densely populated with <u>ICC</u> cells, which generate the “slow wave” electrical activity responsible for initiating the phasic contractions characteristic of this region. These contractions occur in a coordinated process known as peristalsis, propelling food toward the pylorus [24, 19, 23, 22].

Phasic contractions of gastric smooth muscle are driven by neural inputs from the vagal nerve and the <u>ENS,</u> which target the <u>ICC</u> and <u>SMC</u> within the gastric wall [2, 69, 70]. As demonstrated in the study by Athavale et al., 2024 [69], these contractions are regulated by both cholinergic (excitatory) and <u>NANC</u> (inhibitory) pathways. The cholinergic pathway enhances contractile behavior, whereas inhibitory mechanisms—subdivided into purinergic and nitrergic pathways—reduce contraction activity. Together, these pathways modulate the balance of excitatory and inhibitory inputs, finely tuning gastric motility within the antrum [69, 6, 2, 71, 72].

For modeling the motor neuron firing frequency of both excitatory and inhibitory pathways, the <u>MMEHC</u> equation is employed, following a similar methodology as discussed in Section 2.2.1. This modeling approach is integrated into the compartmental framework [19], extending the <u>modified ‘leaky integrate and fire’ (MLIF)</u> model by incorporating excitatory and inhibitory neural inputs modeled by the <u>MMEHC</u> equation for the antrum region (*w* = 2) of the stomach. A schematic representation of this framework is provided in Fig. 2.

**Figure 2.**
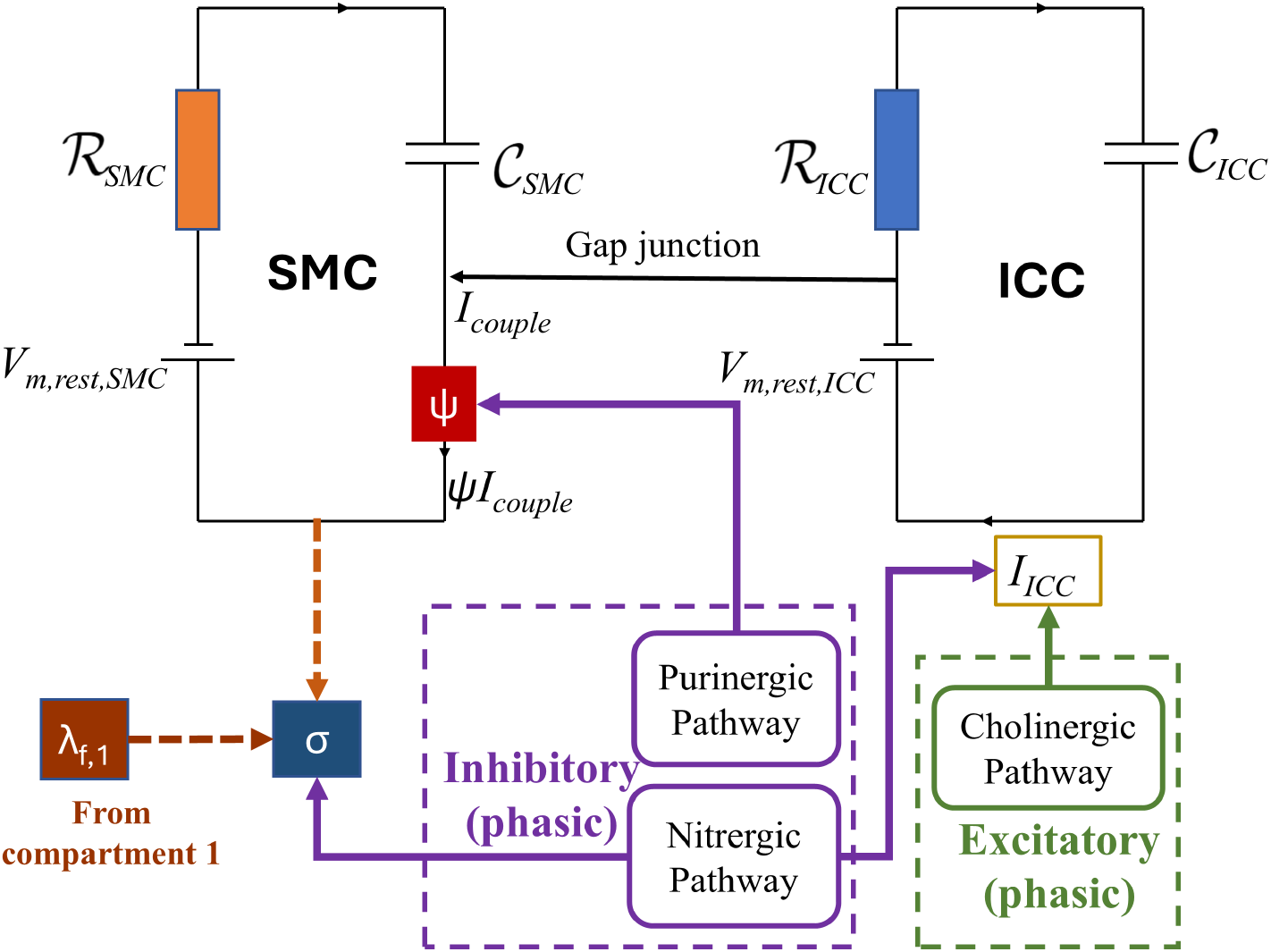
Schematic representation of excitatory and inhibitory stimulation for antrum phasic activity

#### 2.2.6. Cholinergic pathway (Ach): Antrum

In the antrum, the cholinergic pathway is regulated by the neurotransmitter <u>Ach,</u> which is released at the neuromuscular junction. The <u>ICC</u> responds to this neurotransmitter, eliciting an excitatory effect on both the amplitude and frequency of the <u>ICC</u> phasic “slow waves,” which are essential for coordinated motility [69, 73].

The <u>MMEHC</u> equation is used to model the release of acetylcholine [*Ach*]_2_ in response to the firing frequency of the cholinergic pathway *f*_*e*,*p*,2_. The equation is expressed as

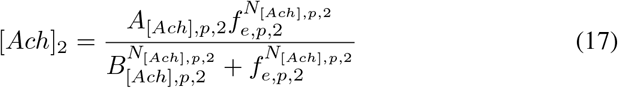

The fractional increase in the <u>ICC</u> “slow wave” amplitude, *ϕ*_*ea*,2_, in response to the acetylcholine concentration [*Ach*]_2_, is modeled using the following equation

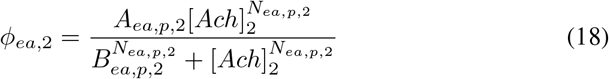

The fractional increase of <u>ICC</u> “slow wave” frequency, *ϕ*_*ca*,2_, in response to acetylcholine concentration is expressed using the following equation

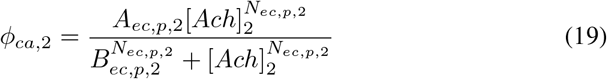

Data from studies by Sinn et al., 2010 and Forrest et al., 2006 [74, 71] was utilized to fit the parameters of the <u>MMEHC</u> equations describing the relationship between cholinergic pathway firing frequency and <u>ICC</u> “slow wave” frequency. Additionally, data from Athavale et al., 2024 [69] and Nakamura et al., 2004 [73] was employed to determine the parameters of the <u>MMEHC</u> equations modeling the increase in <u>ICC</u> “slow wave” amplitude as a response to cholinergic pathway firing frequency.

#### 2.2.7. NANC pathway (NO and purinergic): Antrum

The inhibitory response in the antrum is modulated by the <u>NANC</u> pathway, which includes nitrergic neurotransmitters like <u>NO</u> and purinergic neurotransmitters like <u>adenosine triphosphate (ATP),</u> released at the neuromuscular junction [69, 75, 76]. These neurotransmitters act on smooth muscle and interstitial cells to decrease the amplitude and frequency of contractions, playing a vital role in balancing excitatory and inhibitory inputs within the gastric motility system.

To model nitrergic neurotransmitter release in <u>ICC,</u> the concentration of nitric oxide <u>(NO)</u> neurotransmitter, [*NO*]_*p*,2_, at the neuromuscular junction is governed by the <u>MMEHC</u> equation. This release is modulated by the firing frequency of the <u>NANC</u> pathway, *f*_*i*,*p*,2_, which influences inhibitory signaling and interstitial cell activity. The equation is formulated as follows:

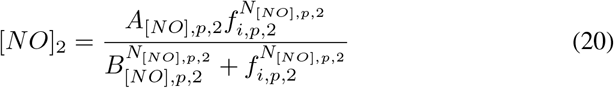

The fractional reduction in <u>ICC</u> “slow wave” amplitude, *ϕ*_*ia*,2_, based on the <u>NO</u> concentration, is represented using the <u>MMEHC</u> equation as follows

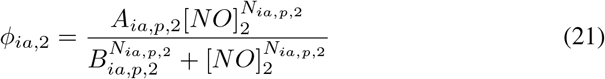

The fractional reduction in <u>ICC</u> “slow wave” frequency, *ϕ*_*ic*,2_, based on the <u>NO</u> concentration, is denoted by the following equation

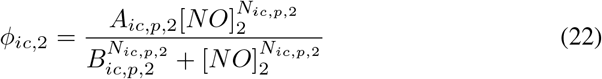

To model purinergic neurotransmitter release in <u>SMC,</u> the concentration of purinergic neurotransmitter [*Pur*]_2_ at the neuromuscular junction is governed by the <u>MMEHC</u> equation. This release is regulated by the firing frequency of the efferent neuron, *f*_*i*,*p*,2_, which influences neurotransmitter availability and subsequent smooth muscle response. The equation is formulated as follows

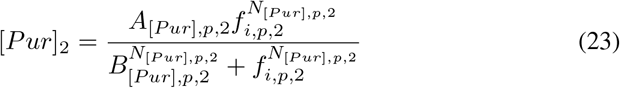

The purinergic neurotransmitter at the neuromuscular junction modulates the <u>SMC</u> “slow wave” amplitude [69]. The fractional reduction in the <u>SMC</u> “slow wave” amplitude, denoted as *ϕ*_*iψ*,2_, in response to the purinergic neurotransmitter concentration is described by the <u>MMEHC</u> equation as follows

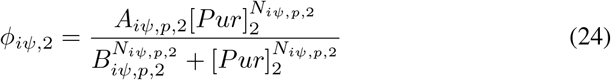

The inhibitory fractional response to the purinergic neurotransmitter, denoted as *ψ*_2_, is represented by the following equation

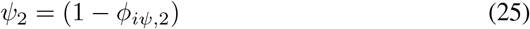

#### 2.2.8. Equations in the compartmental model for the gastric antrum

The equation for the <u>ICC</u> active stimulating current amplitude, which incorporates both the fractional excitatory and inhibitory responses, is denoted as follows

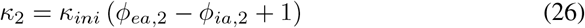

Here, *κ*_*ini*_ represents the baseline active stimulating current amplitude. The equation for the <u>ICC</u> “slow wave” frequency, which incorporates both the fractional excitatory and inhibitory responses, is denoted as follows

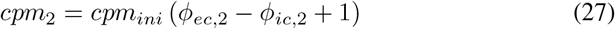

Here, *cpm*_*ini*_ represents the baseline <u>ICC</u> “slow wave” frequency.

The equation for the parameter in the <u>MLIF</u> model that influences the time for each wave cycle, *t*_*end*,2_, is denoted as follows

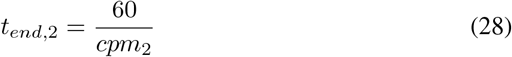

More details on *κ*_*m*_ and *t*_*end*,2_ can be found in our previous study [19].

For the <u>ICC,</u> the “slow wave” activity is represented by the <u>MLIF</u> model [19] as

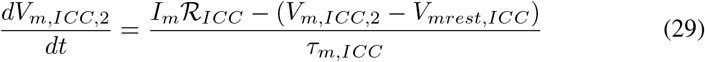

Where

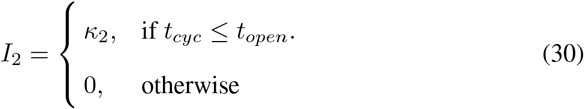

A gap junction equation connects the <u>ICC</u> and <u>SMC</u> models which is denoted by

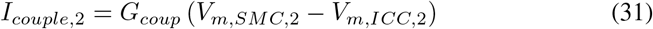

For the <u>SMC</u> “slow wave” activity is modeled [19] as

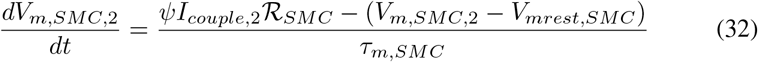

The translation of <u>SMC</u> membrane voltage, *V*_*m*,*SMC*,2_, to activated <u>MLCK,</u> [*MLCK*]_*p*,2_, is based on the framework introduced in our previous work [19]. This framework builds upon models developed by Corrias and Buist, 2007 [77] and Gajendran and Buist, 2011 [28].

The total <u>MLCP</u> concentration, [*MLCP*]_*p*,2_, is expressed as [*MLCP*]_*p*,2_ = [*MLCP*]_*rest*_+ [*MLCP*]_*f*,*p*,2_ where [*MLCP*]_*f*,*p*,2_ is computed using Eqs. <u>8</u> and <u>9.</u>

The interplay between [*MLCP*]_*p*,2_ and [*MLCK*]_*p*,2_ regulates the number of total cross-bridges formed ([*AM*_*p*_]_*p*,2_ + [*AM*]_*p*,2_), which directly influence active muscle contractions [78, 79]. The formation of cross-bridges is modeled using the Hai-Murphy, 1988 framework [65], as represented by Eqs. 10, 11, 12, and 13.

The tissue stress, *σ*_2_, in the antrum is influenced by sarcomere deformation in the fundus compartment, denoted as *λ*_*f*,1_. This deformation reflects the passive distension of the stomach, which may result from the gastric volume of liquid and the total number of cross-bridges formed [31]. The muscle tissue stress *σ*_2_ is modeled using a framework established in previous studies [80, 81, 31]. The governing equations for this relationship are as follows

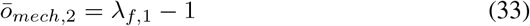

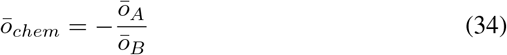

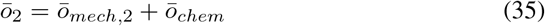

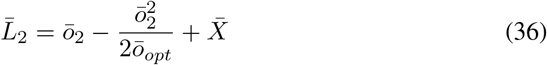

Here, *ō*_*mech*,2_ represents the sliding filament component caused by passive deformation, while *ō*_*chem*,2_ corresponds to the sliding element resulting from active cross-bridging. The normalized overlap of actin-myosin filament lengths is denoted as 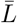. Since the stomach empties gradually rather than instantaneously, the components controlling the stress in the stomach tissue are assumed to be in a steady state. This assumption ensures that the internal and active stresses remain equal at all times. Under this condition, *σ*_2_ is expressed as follows [80, 31]

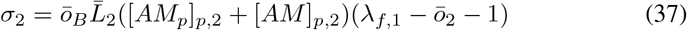

The <u>non-linear viscoelastic model (NLVM)</u> model is used to translate tissue stress into tissue stretch, following a framework similar to that of our previous study [19]. However, for the principal stress *E* of the hyperelastic tissue model, a polynomial equation is employed to represent the hyperelastic material, as demonstrated in the study by Panda and Buist, 2018 [82]. For further details, refer to Section 8.2.2. The principal stress *E*_2_ is expressed as follows

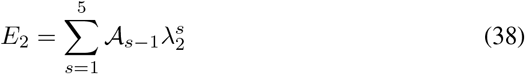

The viscoelastic property, or hysteresis loop, is captured using a nonlinear dashpot model, *η*_2,2_, similar to the approach described in our previous work [19]. The tissue stretch *λ*_2_ equation resulting from tissue stress is expressed as

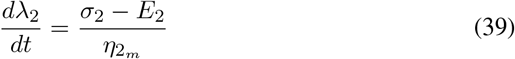

Where

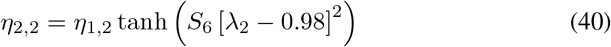

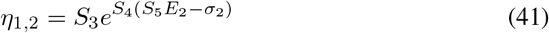

The radius of the cylindrical compartment, *r*_*fin*,2_, modeled using the framework from our previous study [19], is expressed as

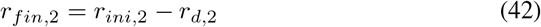

where

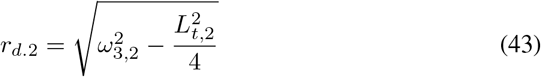

Here, *r*_*d*,2_ represents the deformed compartmental radius, *L*_*t*,2_ denotes the undeformed tissue length, and *ω*_3,2_ is the component that accounts for the deformed tissue length.

#### 2.2.9. Motor neurons influencing tonic and phasic activity: PS

The <u>PS</u> (*w* = 3) regulates gastric flow from the stomach to the duodenum. Vagal regulation of the <u>PS</u> involves both cholinergic and <u>NANC</u> pathways, which are primarily responsible for maintaining the basal tone of the sphincter through tonic contractions [83]. The cholinergic pathway increases basal tone by inducing smooth muscle contraction, while the <u>NANC</u> pathway reduces basal tone by promoting smooth muscle relaxation. <u>Ach</u> is likely the primary neurotransmitter for smooth muscle contraction in the sphincter. However, the complete identity of the inhibitory neurotransmitters involved in smooth muscle relaxation through the <u>NANC</u> pathway remains unclear in the literature [83].

In addition to tonic contractions, the <u>PS</u> exhibits phasic contractions regulated by <u>ICC</u> cells, which share similar properties with those in the antrum of the stomach [84, 85, 42]. Like the antrum and fundus, the <u>PS</u> comprises circular and longitudinal muscle layers that exhibit elastic properties for storing passive stress [86]. Changes in basal tone can induce passive stress, which, in turn, may influence the active stress generated by the contractile elements in the muscle.

For modeling purposes, the signaling framework developed for phasic contractions in the antrum is adapted for the <u>PS.</u> For tonic activity, separate pathways for cholinergic and <u>NANC</u> signaling will be derived. The schematic for the phasic and tonic activity in the <u>PS</u> is shown in Fig. 3.

**Figure 3.**
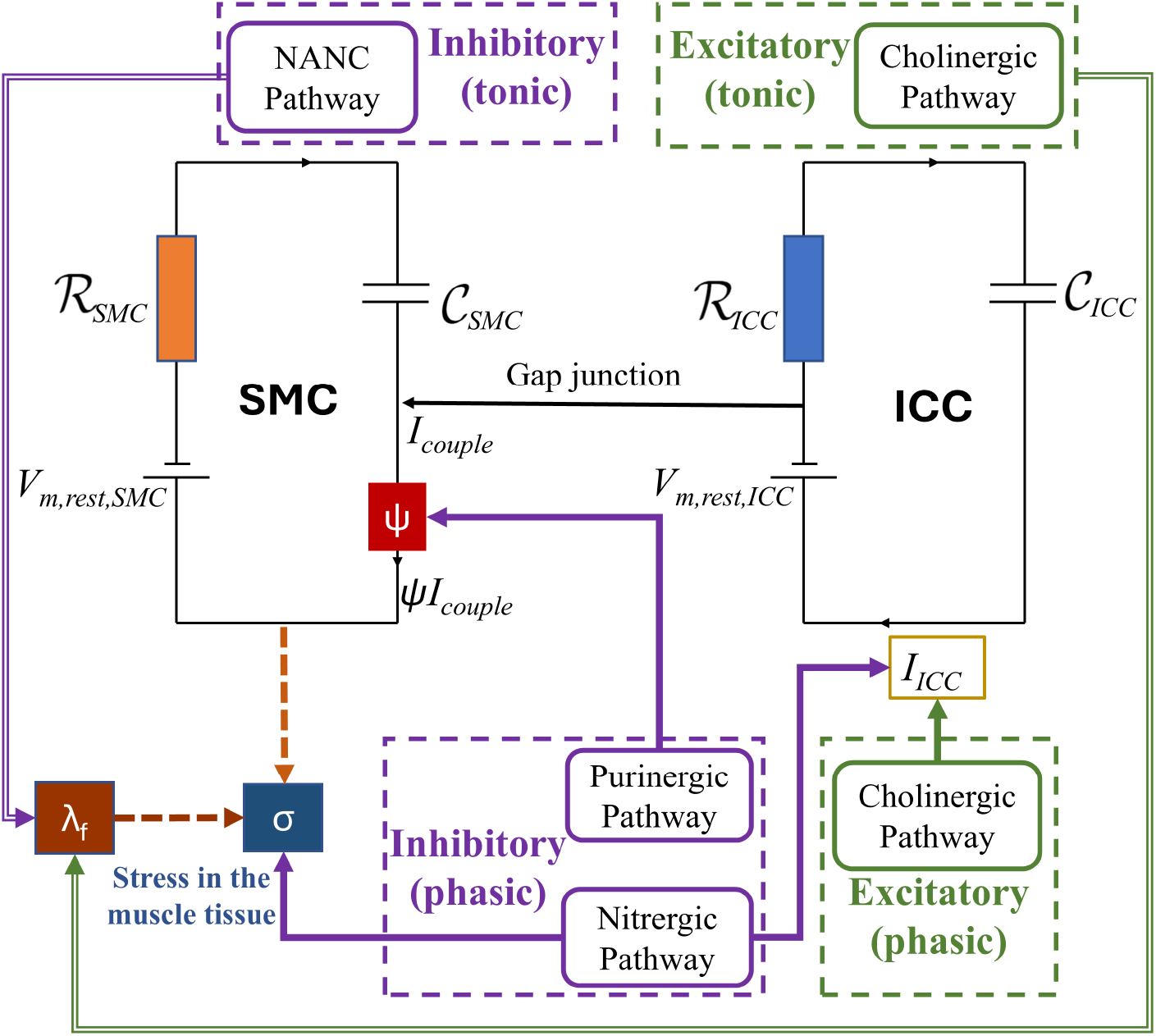
Schematic representation of excitatory and inhibitory stimulation for PS phasic and tonic activity

#### 2.2.10. Cholinergic pathway: PS

For tonic-controlled activity, due to the lack of data in the literature regarding the effect of the cholinergic pathway on <u>PS</u> contractions, the modeling approach for this pathway is based on equations derived for tonic activity in the fundus. These equations are represented as Eqs. 2 and 3, where *f*_*e*,*to*,3_ denotes the excitatory firing frequency that regulates tonic activity.

For phasic-controlled activity, the cholinergic pathway neuron is modeled similarly to that of the antrum, as the <u>ICC</u> cells in the <u>PS</u> exhibit properties comparable to those in the antrum. Here, the excitatory firing frequency governing phasic activity is represented as *f*_*e*,*p*,3_.

#### 2.2.11. NANC pathway: PS

For tonic-controlled activity, the identity of the neurotransmitter responsible for inhibitory activity is unknown [83]. Therefore, the <u>MMEHC</u> equation is used directly to model the inhibitory response, where *f*_*i*,*to*,3_ represents the inhibitory firing frequency, and *ϕ*_*ih*,*to*,3_ denotes the inhibitory fraction. The equation is expressed as

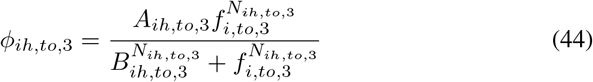

Data from Ishiguchi et al., 2000 [87] was used to fit the parameters for the equation above.

For phasic activity, the <u>NANC</u> pathway is modeled similarly to that of the antrum, where the inhibitory firing frequency is represented as *f*_*i*,*p*,3_.

#### 2.2.12. Equations in the compartmental model for PS

For tonic activity, the intracellular calcium concentration in the <u>SMC</u> is modeled using Eq. 7. The model by Gajendiran and Buist, 2011 [28] is employed to convert 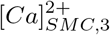 into active <u>MLCK</u>, denoted as [*MLCK*_*act*_]_*to*,3_.

Due to the fact that the neurotransmitter involved in the <u>NANC</u> pathway is unknown, the pathway is modeled such that *ϕ*_*ih*,*to*,3_ increases <u>MLCP</u> and decreases <u>MLCK</u> concentrations, reflecting effects observed in inhibitory pathways [29]. The <u>MLCK</u> and <u>MLCP</u> concentrations associated with the inhibitory pathway are modeled using the following equations as [*MLCP*]_*to*,3_ = [*MLCP*]_*rest*_+*ϕ*_*ih*,*to*,3_[*MLCP*]_*f*,*to*,3_ and [*MLCK*]_*to*,3_ = (1 − *ϕ*_*ih*,*to*,3_) [*MLCK*_*act*_]_*to*,_.

To model the effect of <u>MLCP</u> and <u>MLCK</u> pathway concentrations on tissue deformation due to passive stress (tonic activity), Eqs. 10 to 16 are utilized, with detailed explanations provided in Section 2.2.4.

For phasic activity in the <u>PS,</u> the effects of the cholinergic and <u>NANC</u> pathways on the <u>ICC, SMC,</u> and tissue deformation are modeled using equations similar to those developed for the antrum. The details of the model are explained in Section 2.2.8.

To determine the gastric flow rate through the <u>PS,</u> *Q*_*flow*_, a simplified flow rate equation is used

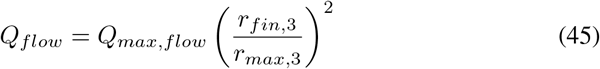

where *Q*_*max*,*flow*_ represents the maximum flow rate through the <u>PS,</u> and *r*_*max*,3_ is the maximum radius the <u>PS</u> can achieve. For further details, refer to Section 8.2.3.

### 2.3. Modeling the afferent/sensory neuron connections

The ascending vagal nerves, also known as vagal afferent fibers, constitute approximately 80 % of all vagal fibers. These sensory neurons detect local chemical and mechanical signals from the <u>GI</u> tract and relay this information to the brainstem for processing. There are two primary categories of vagal afferents: chemosensitive and mechanosensitive (1) Chemosensitive fibers have peripheral endings that respond to chemical stimuli, such as nutrient content, pH levels, hormones, and immune signals. (2) Mechanosensitive afferents include mucosal endings, <u>intramuscular arrays (IMAs),</u> and <u>intraganglionic laminar endings (IGLEs),</u> which respond to mechanical stimuli like mucosal stroking, muscle distension, and contraction [88, 89].

The stomach is rich in mechanosensitive sensory neurons, including mucosal endings, <u>IGLEs,</u> and <u>IMAs,</u> making it highly responsive to stretch and contractions. When the stomach wall distends due to increased gastric volume, mechanosensitive afferent fibers demonstrate higher firing frequencies. This relationship was evidenced in a study by Williams et al., 2016 [90].

In contrast, chemosensitive afferents are abundant in the proximal small intestine, where they detect nutrient content and signal the brainstem to modulate gastric tone. For instance, chemosensitive signals can increase the basal tone of the <u>PS</u> to slow gastric emptying, allowing sufficient time for nutrient digestion in the small intestine [83, 91]. This study excludes pH-sensitive chemosensitive afferents since the compartmental model framework does not incorporate microbiome interactions or chemical reactions affecting gut pH.

From a compartmental modeling perspective, two afferent pathways are considered: (a) a mechanosensitive pathway detecting muscle length stretch, influenced by stomach volume changes; and (b) a chemosensitive pathway sensing the nutrient content flow rate at the <u>PS</u> as gastric contents flow into the proximal duodenum.

The firing rates of mechanosensitive and chemosensitive receptors are modeled using linear equations. Linear models are appropriate because muscle stretch and nutrient flow rate changes gradually rather than abruptly. Studies [92, 93] have shown that non-linear models are only necessary for abrupt sensory responses. For instance, Zagorodnyuk and Brookes, 2000 [92], demonstrated a linear relationship between firing rate and tissue stretch under gradual changes, and Mei, 1977 [94], reported a roughly linear relationship between glucose density and vagal afferent firing.

The mechanoreceptor afferent firing rate *f*_*mech*_, is modeled as

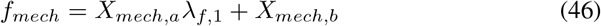

where *X*_*mech*,*a*_ and *X*_*mech*,*b*_ are the mechanosensitive fitting constants,fitted using data from Zagorodnyuk and Brookes, 2000 [92].

For chemoreceptors, nutrient content volumetric flow rate *Q*_*cal*_, is based on Kong and Singh, 2008 [27], which reports typical flow rates of 2–4 kcal/min during gastric emptying. The maximum calorie flow rate per minute is denoted as *γ*_*max*_. A dimen-sionless constant Ω accounts for flow lost during the periodic opening and closing of the <u>PS.</u> The calorie content per unit volume of gastric liquid is represented as *g*_*cal*_. Thus, *Q*_*cal*_ is defined as

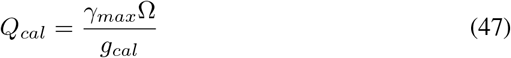

The chemoreceptor afferent firing rate *f*_*chem*_ is modeled by the equation

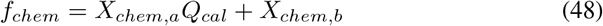

where *X*_*chem*,*a*_ and *X*_*chem*,*b*_ are the chemosensitive fitting constants translating sensory activity to afferent firing rates.

The constants *X*_*chem*,*a*_ and *X*_*chem*,*b*_ were fitted to ensure that the maximum afferent firing rate was 30 Hz, a value reported by Mei, 1977 [94]. The parameter Ω was determined by analyzing the gastric emptying rate from a dataset in the study by Kwiatek et al., 2009 [95].

Additionally, the model incorporates an afferent firing response triggered by “fight or flight” conditions. This response is modeled as an on-off (binary) variable 𝒪_*sym*_, where 𝒪_*sym*_ = 1 indicates the presence and 𝒪_*sym*_ = 0 indicates the absence of a sympathetic response. When activated, the sympathetic response overrides all parasympathetic pathways, inhibiting gastric emptying by reducing peristaltic activity, maintaining pyloric sphincter closure.

### 2.4. Modeling the interneuron connection in the brainstem

The neurochemically and biophysically diverse second-order neurons of the <u>NTS</u> process sensory information transmitted by vagal neurons. Vagal afferent fibers carry mechanical, chemical, and osmotic signals from the viscera to the <u>NTS,</u> where this information integrates with brainstem, limbic, and hypothalamic inputs to ensure optimal control of stomach reflexes, motility, and emptying [2, 6, 96, 11, 12].

Glutamate is the primary neurotransmitter used by all vagal afferents, irrespective of their modality or function, to relay information to the <u>NTS.</u> Activation of sensory vagal afferent pathways triggers second-order <u>NTS</u> neurons via glutamate action on <u>N-methyl D-aspartate (NMDA)</u> and <u>non-NMDA</u> receptors, initiating reflex activities. These second-order neurons utilize various neurotransmitters to regulate the output of <u>DMV</u> neurons, which govern gastric functions and close the vago-vagal reflex loop [2]. The topographic organization of visceral sensory afferents within <u>NTS</u> subnuclei introduces spatial heterogeneity in how sensory information is processed and relayed to the brainstem [2].

The <u>NTS</u> provides key synaptic inputs to <u>DMV</u> neurons, which play a critical role in controlling vago-vagal responses. Among these inputs, GABAergic projections are central in modulating <u>DMV</u> neuronal firing rates, thereby influencing vagal efferent output that regulates gastric tone and motility. Blocking GABAergic transmission between the <u>NTS</u> and <u>DMV</u> using the <u>gamma-aminobutyric acid (GABA)</u>_A_ antagonist bicuculline has been shown to increase the firing rate of most <u>DMV</u> neurons, resulting in enhanced gastric motility and tone [6].

The vagal efferent or motor inputs modulated by <u>DMV</u> neurons are controlled through cholinergic and <u>NANC</u> pathways. The interneuronal connections between the <u>NTS</u> and <u>DMV,</u> whether direct or indirect, link the afferent sensory neuron firing frequency to the efferent motor neuron firing frequency. This relationship is modeled using a sigmoid curve equation, as previously demonstrated by Park et al., 2020 [97].

Park et al., 2020 [97], modeled the brainstem afferent-efferent firing frequency relationship in the cardiovascular system. Similar plasticity is observed in brainstem interactions within the <u>GI</u> system, where motor neuron firing adapts and fine-tunes in response to sensory neuron activity [97, 98, 99].

In this study, the equation derived by Park et al., 2020 [97] is utilized to model the relationship between vagal afferent and vagal efferent mechanosensitive responses. This equation models the passive tissue stretch, caused by stomach volume changes, to the efferent firing rate driving peristaltic activity in the antrum. A detailed explanation of this pathway is provided in Section <u>2.5.</u> The mechanosensitive afferent-efferent relationship is described by the equation

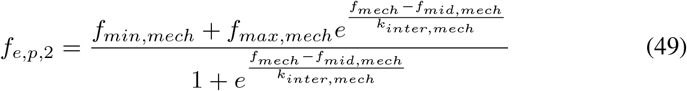

For the chemosensitive response, which relates nutrient content flow rate to <u>PS</u> opening, a ninth-order polynomial equation is employed. This choice is based on its superior fit for modeling afferent-efferent responses compared to the equation from Park et al., 2020 [97]. A comprehensive understanding of this pathway is also available in Section <u>2.5.</u> The polynomial equation is expressed as

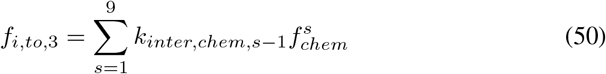

For more details on the interneuron brainstem connection equations and their corresponding parameters, refer to Section 8.3.

### 2.5. Gut-brain axis: Sympathetic and parasympathetic regulation of gastric function

The gut–brain axis integrates both sympathetic and parasympathetic pathways, as illustrated in Fig. 4 A). The sympathetic branch is typically activated during the “fight or flight” response, where evidence has shown that digestive activity is suppressed [100, 10, 6]. In this study, the model simulates inhibition of the digestion process by restricting gastric motility.

**Figure 4.**
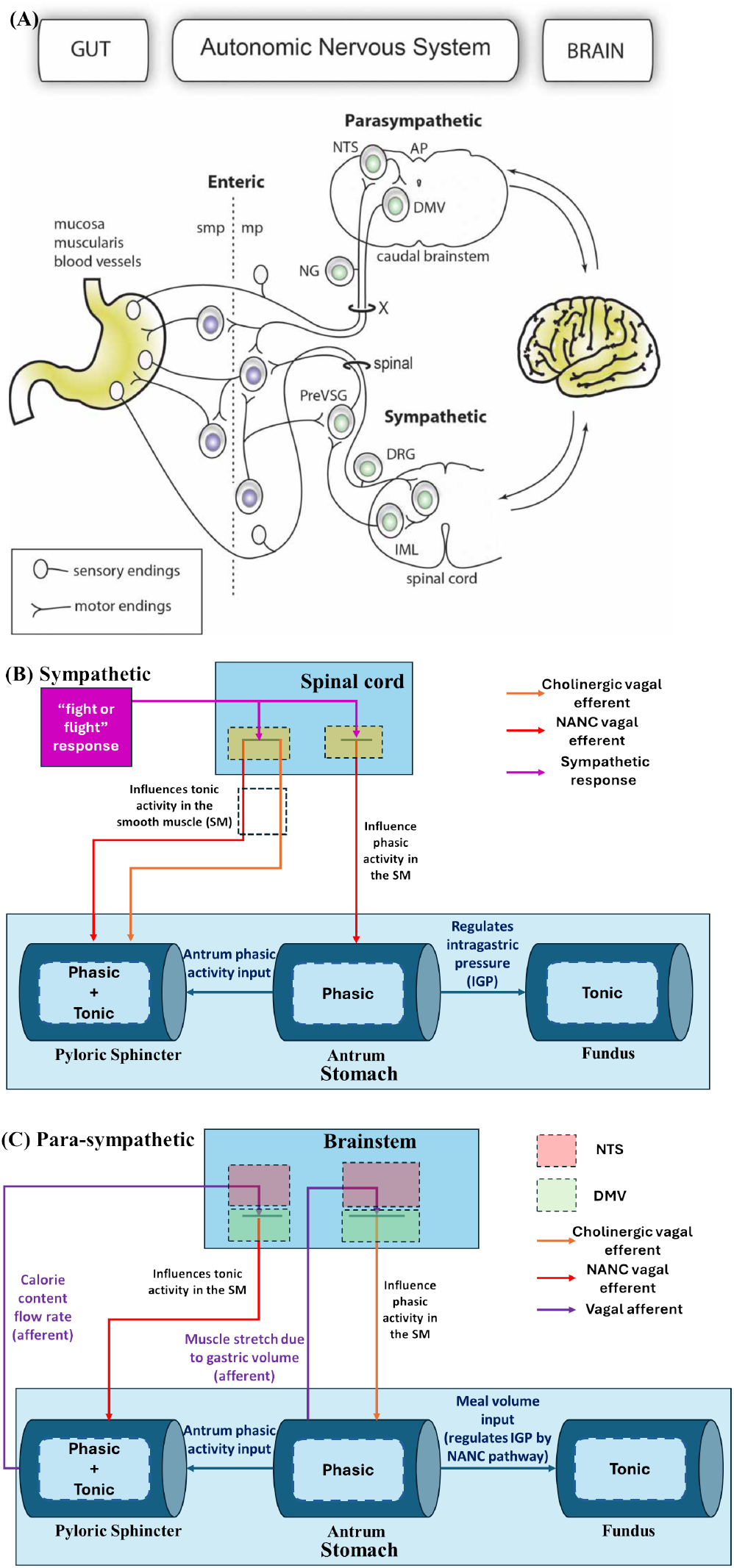
A) Gut-brain axis schematic: sympathetic and para-sympathetic pathways for gastric function regulation by Udit and Gautron, 2013 [100] B) Compartmental model schematic for sympathetic pathway C) Compartmental model schematic for parasympathetic pathway for gastric function regulation during gastric emptying

In the antrum, phasic activity is inhibited through the <u>NANC</u> pathway, which suppresses peristaltic activity. In the <u>PS,</u> the frequency of the <u>NANC</u> pathway is reduced, while the cholinergic pathway is upregulated, leading to the closure of the <u>PS.</u> This prevents gastric liquid flow from the stomach to the duodenum. Upon activation of the sympathetic response, the efferent neuron frequencies *f*_*i*,*p*,2_ and *f*_*e*,*p*,2_ in the antrum are set to 15 Hz and 0 Hz, respectively. Similarly, the frequencies *f*_*i*,*to*,3_ and *f*_*e*,*to*,3_ in the <u>PS</u> are set to 0 Hz and 10 Hz, respectively. These motor neuron efferent frequencies result in inhibitory gastric emptying by closing the <u>PS</u> and reducing peristaltic activity. The <u>intragastric pressure (IGP)</u> is maintained in the fundus compartment via intramural firing of the <u>NANC</u> pathway, which adjusts gastric volume. A schematic representation of the model for the sympathetic pathway is shown in Fig. 4 B).

Since this study focuses on the gut-brain axis during the gastric emptying phase, the parasympathetic pathway is modeled to regulate gastric functions during this phase. The fundus compartment is connected intramurally via the <u>NANC</u> pathway, which is active during gastric emptying to facilitate adaptive relaxation (or gastric accommodation) [46].

A piecewise polynomial equation is used to model the intramural connection of the fundus, establishing the relationship between stomach volume *V*_*tot*_ and the firing frequency of the fundic <u>NANC</u> pathway neurons in the <u>ENS,</u> which plays a crucial role in gastric relaxation. Further details on the use of the piecewise polynomial equation and its fitting parameters to model the intramural connection are provided in Section 8.3. The piecewise polynomial equation is expressed as

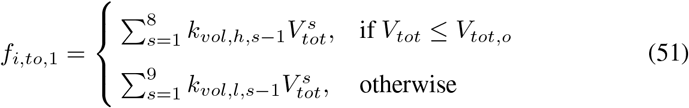

If the stomach is overly full, stronger peristaltic contractions in the antrum are required, which can be achieved by activating the cholinergic pathway. For gastric emptying, the <u>PS</u> must relax to allow gastric liquid to flow from the stomach to the duodenum. This relaxation is mediated by the <u>NANC</u> pathway. Sensory pathways, both mechanoreceptive and chemoreceptive, relay information about stomach tissue stretch to determine fullness and monitor the gastric meal flow rate from the stomach to the duodenum.

Phasic contractions of the <u>PS,</u> which open and close periodically (approximately 3 <u>cycles per minute (cpm)) [101],</u> are primarily controlled intramurally. However, they may also receive vagal inputs [45]. These phasic contractions, driven by the <u>ICC,</u> close the <u>PS</u> in response to peristaltic waves reaching the terminal antrum [19]. For simplicity, and due to limited evidence of vagal control over phasic <u>PS</u> contractions, the proposed model assumes that <u>ICC-re</u>gulated phasic contractions occur at baseline levels, independent of calorie content. Instead, these phasic contractions are regulated by antral contractions of the stomach.

The schematic representation of the parasympathetic pathway used in this study is illustrated in Fig. 4 C).

## 3. Results and discussion

The compartmental model in this study was solved using MATLAB R2024b and consisted of approximately 70 <u>ordinary differential equations (ODEs)</u> and 81 algebraic equations. Various <u>ordinary differential equation (ODE)</u> solvers in MATLAB, such as ‘ode15s’, ‘ode23s’, ‘ode23t’, etc., were capable of solving the model on a standard home laptop. Among them, the ‘ode15s’ solver demonstrated the most efficient computational performance. The computational time was within a single order of magnitude of seconds, for simulating approximately 200 seconds of model dynamics.

### 3.1. Parasympathetic and sympathetic response

To compute the parasympathetic and sympathetic responses, the total gastric volume, *V*_*tot*_, was set to 0.6 L for these simulations. For the parasympathetic response, the 𝒪_*sym*_ value was set to 0, whereas for the sympathetic response, the 𝒪_*sym*_ value was set to 1. The results obtained from these inputs are shown in Fig. 5.

**Figure 5.**
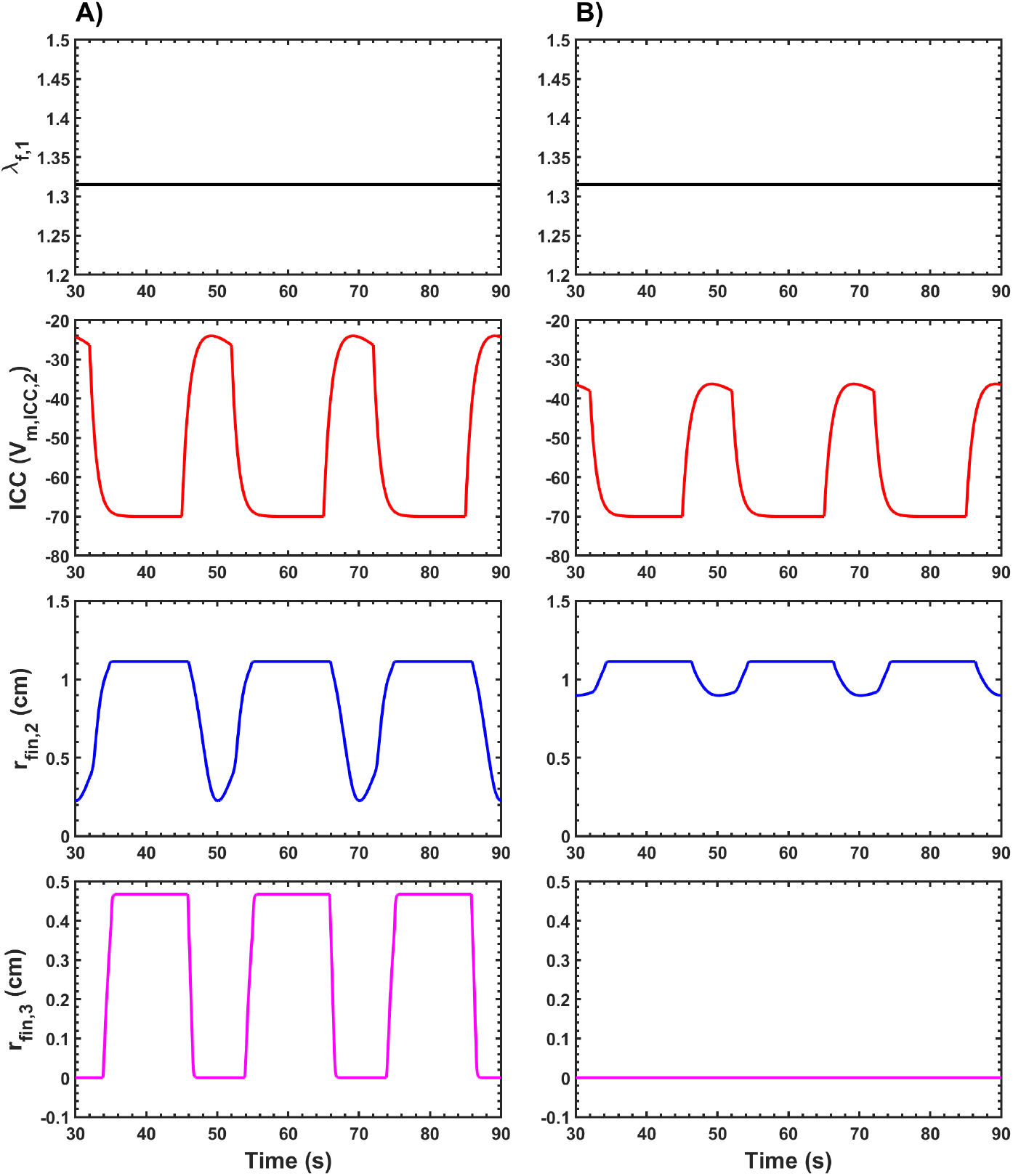
Simulated data showing *λ*_*f*,1_, ICC “slow wave” membrane potential *V*_*m*,*ICC*,2_ (mV) in the antrum compartment, radius of the antrum compartment *r*_*fin*,2_ (cm), and radius of the PS compartment *r*_*fin*,3_ (cm) plotted against time. Results are shown for A) Parasympathetic and B) Sympathetic response

The *λ*_*f*,1_ value, which determines fundic relaxation, remains the same for both the parasympathetic and sympathetic responses. This is because fundic relaxation depends on *V*_*tot*_, which is itself determined by the gastric contents or meal volume and the volume required to maintain <u>IGP</u> in the stomach. Since *V*_*tot*_ is identical for both responses, the *λ*_*f*,1_ value is also identical, ensuring that the fundus can accommodate the meal contents while maintaining <u>IGP.</u>

For the antrum, the <u>ICC</u> activity, *V*_*m*,*ICC*,2_, is reduced during the sympathetic response compared to the parasympathetic response. This reduction in <u>ICC</u> “slow wave” activity affects peristaltic contractions in the antrum, as seen in Fig. 5. During the parasympathetic response, the radius of the antral compartment (*r*_*fin*,2_) decreases significantly, with an occlusion of approximately 79 %. In contrast, during the sympathetic response, the peristaltic contraction is much weaker, with an occlusion of less than 20 %. These findings demonstrate an inhibitory effect on antral peristalsis during the sympathetic response, which aligns with observations reported in the literature [102].

For the <u>PS,</u> Fig. 5 illustrates distinct differences between the parasympathetic and sympathetic responses. During the parasympathetic response, the <u>PS</u> undergoes periodic contractions and relaxations, allowing the sphincter to open and close, thereby facilitating gastric emptying of contents into the small intestine. When relaxed, the radius of the <u>PS</u> is approximately 0.46 cm. However, during the sympathetic response, the <u>PS</u> remains fully contracted, with a radius of 0 cm, preventing gastric contents from entering the small intestine. These results—promoting gastric emptying during the parasympathetic response and inhibiting it during the sympathetic response—are consistent with findings in the literature [103, 104].

### 3.2. Fundus and antrum activity during gastric emptying

To investigate the mechanical behavior of the fundic and antral regions during gastric emptying, simulations were performed at three distinct gastric volumes, *V*_*tot*_: 0.2 L, 0.5 L, and 1.1 L. The corresponding results are presented in Fig. 6. As gastric volume increased, a corresponding rise in the fundic relaxation parameter *λ*_*f*,1_ was observed. Specifically, higher *λ*_*f*,1_ values were associated with greater *V*_*tot*_, indicating enhanced relaxation of the fundus. This response is physiologically consistent with the adaptive relaxation mechanism of the proximal stomach and aligns with experimental findings reported in previous studies [32, 105, 106].

**Figure 6.**
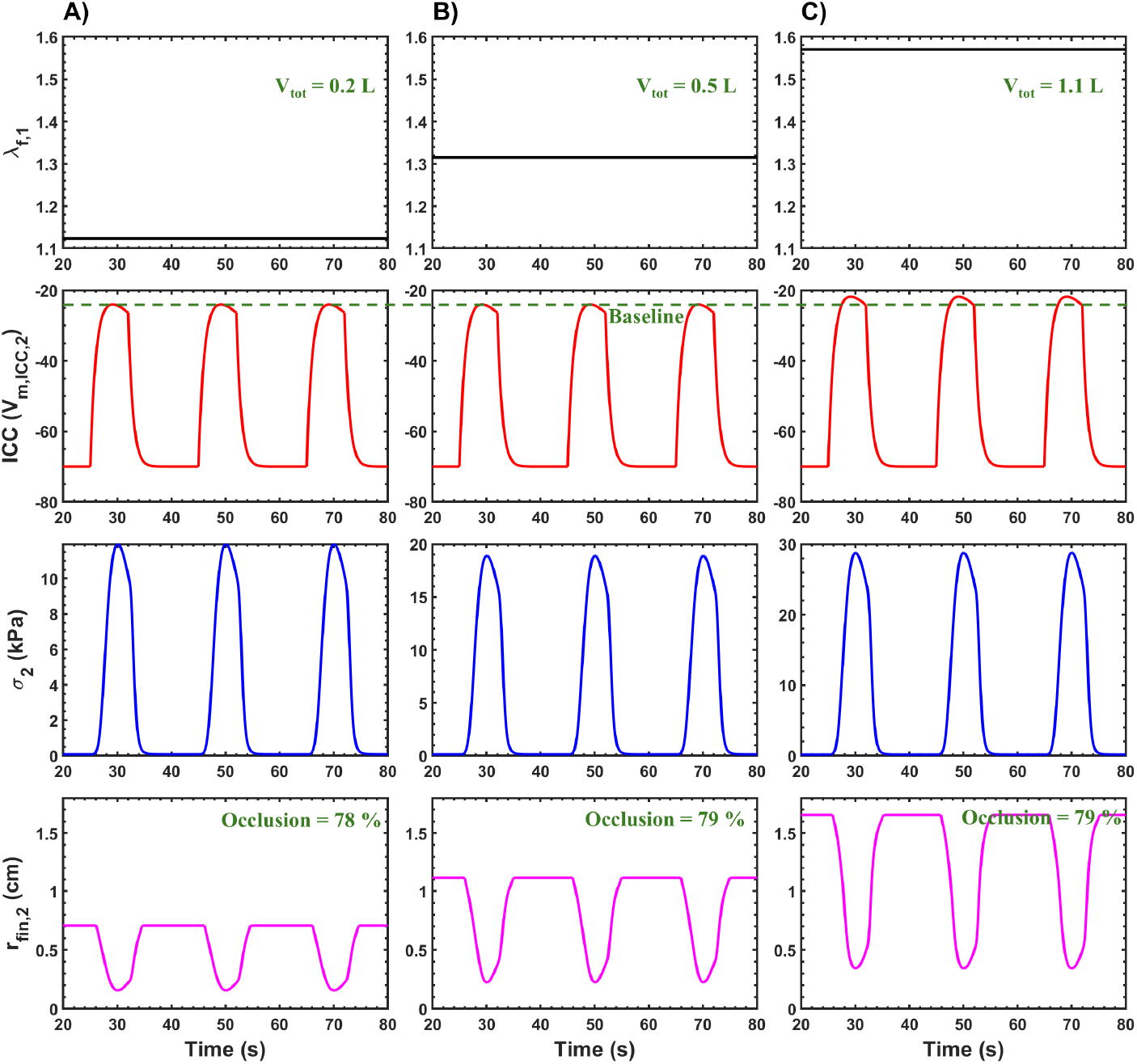
Simulated data showing *λ*_*f*,1_, ICC “slow wave” membrane potential *V*_*m*,*ICC*,2_ (mV) in the antrum compartment, muscle tissue stress in the antrum compartment *σ*_2_ (kPa) and radius of the antrum compartment *r*_*fin*,2_ (cm) plotted against time. Results are presented for total gastric volume *V*_*tot*_ of A) 0.2 L B) 0.5 L and C) 1.1 L

The amplitude of the interstitial cells of Cajal (ICC) “slow wave” membrane potential, *V*_*m*,*ICC*,2_, remained comparable for *V*_*tot*_ = 0.2 L and *V*_*tot*_ = 0.5 L. However, a marked increase in amplitude was observed at *V*_*tot*_ = 1.1 L. This suggests that at higher gastric loads, excitatory signaling to the ICC network is amplified to facilitate stronger contractile activity, likely as a compensatory mechanism to handle the increased luminal contents.

The antral tissue stress, *σ*_2_, exhibited a clear volumetric dependence, with higher *V*_*tot*_ values resulting in increased stress amplitudes (Fig. 6). This is attributed to the bidirectional coupling between electrical and mechanical activity, wherein elevated “slow wave” amplitudes enhance muscle activation. Additionally, passive mechanical stress contributes to sarcomere elongation within the muscle fibers, influencing the active contractile response through length-tension interactions. This integrative electro-mechanical behavior is supported by prior modeling and experimental studies [107, 80, 31, 39].

Despite variations in initial muscle state, the degree of occlusion during antral contractions remained consistent across all volumes, with an average occlusion of approximately 78–79%. However, the relaxed (pre-contraction) radius of the antral compartment increased proportionally with *V*_*tot*_: approximately 0.7 cm for 0.2 L, 1.1 cm for 0.5 L, and 1.65 cm for 1.1 L. These results indicate that both the fundus and antrum undergo volumetric relaxation at higher gastric volumes, consistent with observations reported by Perlas et al. 2009 [108].

To maintain this consistent occlusion percentage, the absolute contraction amplitude also increased with greater *V*_*tot*_. The simulated amplitude of radius reduction during antral contractions was approximately 0.55 cm at 0.2 L, 0.89 cm at 0.5 L, and 1.31 cm at 1.1 L. This pattern supports the hypothesis that increased luminal load necessitates stronger peristaltic contractions for effective gastric emptying. These findings are in agreement with experimental work by Stemper et al. 1975 [109], which reported enhanced antral contractility in response to larger meal volumes.

### 3.3. PS activity during gastric emptying for different calorie content meals

As shown in Fig. 7, the simulation results indicate that increasing the *g*_*cal*_ value from approximately 0 to 0.67 decreases the *λ*_*f*,3_ value. In the model, *λ*_*f*,3_ is responsible for inducing passive stress in the <u>PS</u> muscle tissue, which directly affects the basal tone of the <u>PS.</u> When *g*_*cal*_ increases, the <u>PS</u> transitions to a more contracted state, resulting in an increase in basal tone. Consequently, the radius of the <u>PS</u> compartment (*r*_*fin*,3_) in the stimulated model decreases during the <u>PS</u> open state for higher *g*_*cal*_ values. This reduction in the <u>PS</u> radius restricts the flow of gastric contents from the stomach to the small intestine for gastric liquids with higher caloric content.

**Figure 7.**
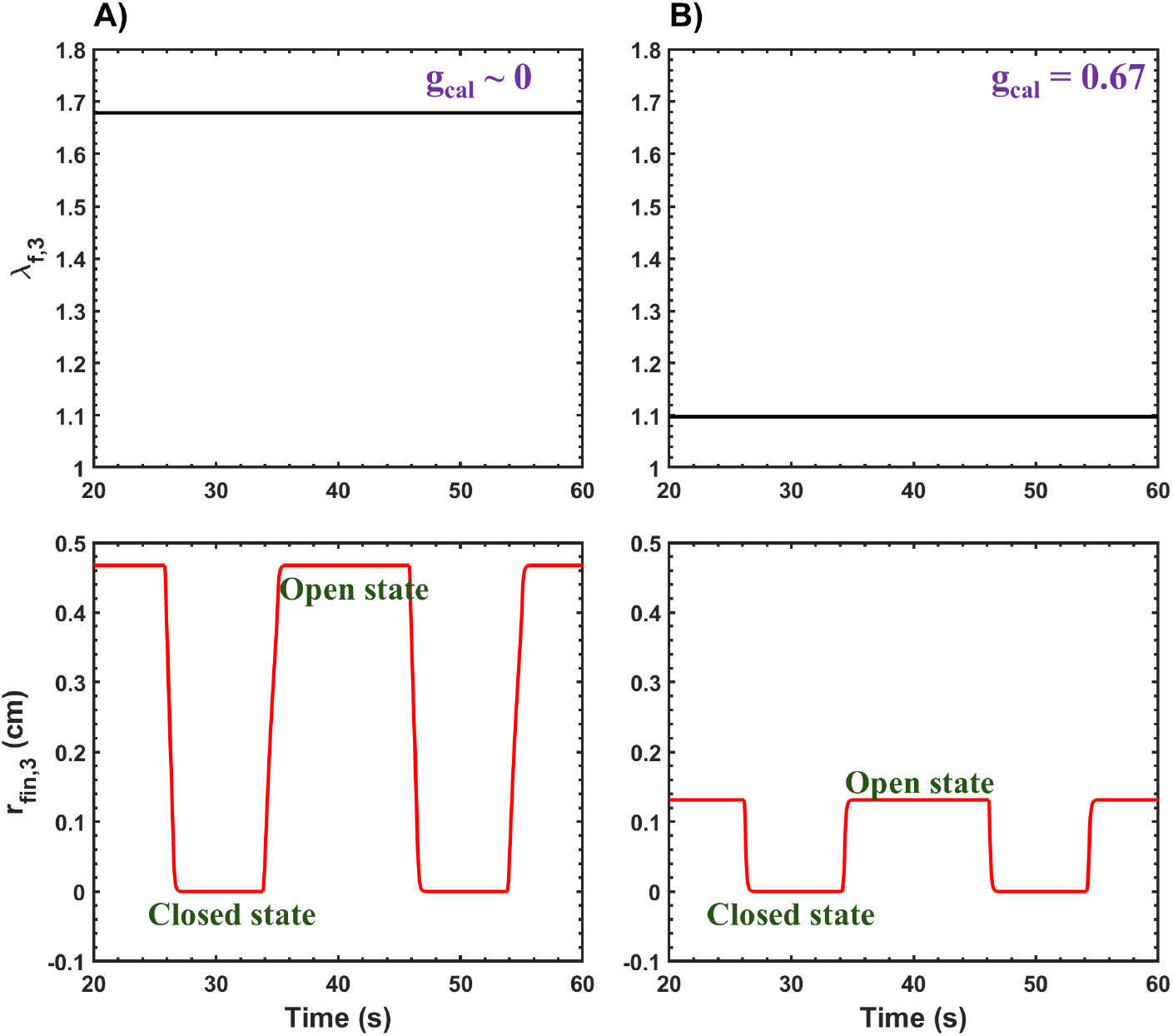
Simulated data showing *λ*_*f*,1_ and the radius of the PS compartment *r*_*fin*,3_ (cm) plotted against time. Results are presented for *g*_*cal*_ values of the gastric liquid: A) approximately 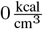 and 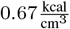

From Fig. 7, the *r*_*fin*,3_ value during the <u>PS</u> open state is approximately 0.46 cm when *g*_*cal*_ is 0, compared to 0.13 cm when *g*_*cal*_ is 0.67. This demonstrates a significant reduction in the <u>PS</u> radius for higher-calorie gastric liquids. These results align with findings in the literature, which report that calorie-dense meals increase levels of <u>cholecystokinin (CCK),</u> subsequently increasing the basal tone of the <u>PS</u> [83, 43, 110, 111, 112].

### 3.4. Impact of meal caloric content on gastric emptying volume

To compute the gastric emptying of a meal, the total gastric volume (*V*_*tot*_) is given by *V*_*tot*_ = *V*_*meal*_ + *V*_*gas*_. Here, *V*_*meal*_ represents the volume of the gastric meal, and *V*_*gas*_ denotes the volume of gas in the stomach, primarily located in the fundus or proximal stomach region [113]. The *V*_*gas*_ is assumed to be constant, as it represents the volume of gas needed to maintain a constant <u>IGP</u> [114], which is calculated using Boyle’s law. Based on a study by Kwiatek et al., 2009 [95], *V*_*gas*_ is approximately 210 mL. The gastric meal emptying is modeled using the equation 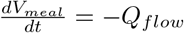, where the initial value of *V*_*meal*_ is the input into the model.

The model computes gastric meal emptying profiles, as shown in Fig. 8. Several simulations were performed and compared with experimental data from Kwiatek et al., 2009 [95], for meals with different caloric densities (*g*_*cal*_) of 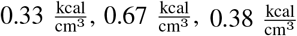, and 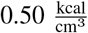. The gastric emptying rates (*Q*_*flow*_) derived from Fig. 8 are approximately 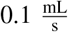 for 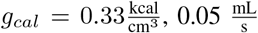 for 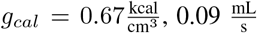 for 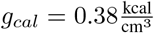, and 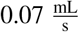 for 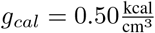.

**Figure 8.**
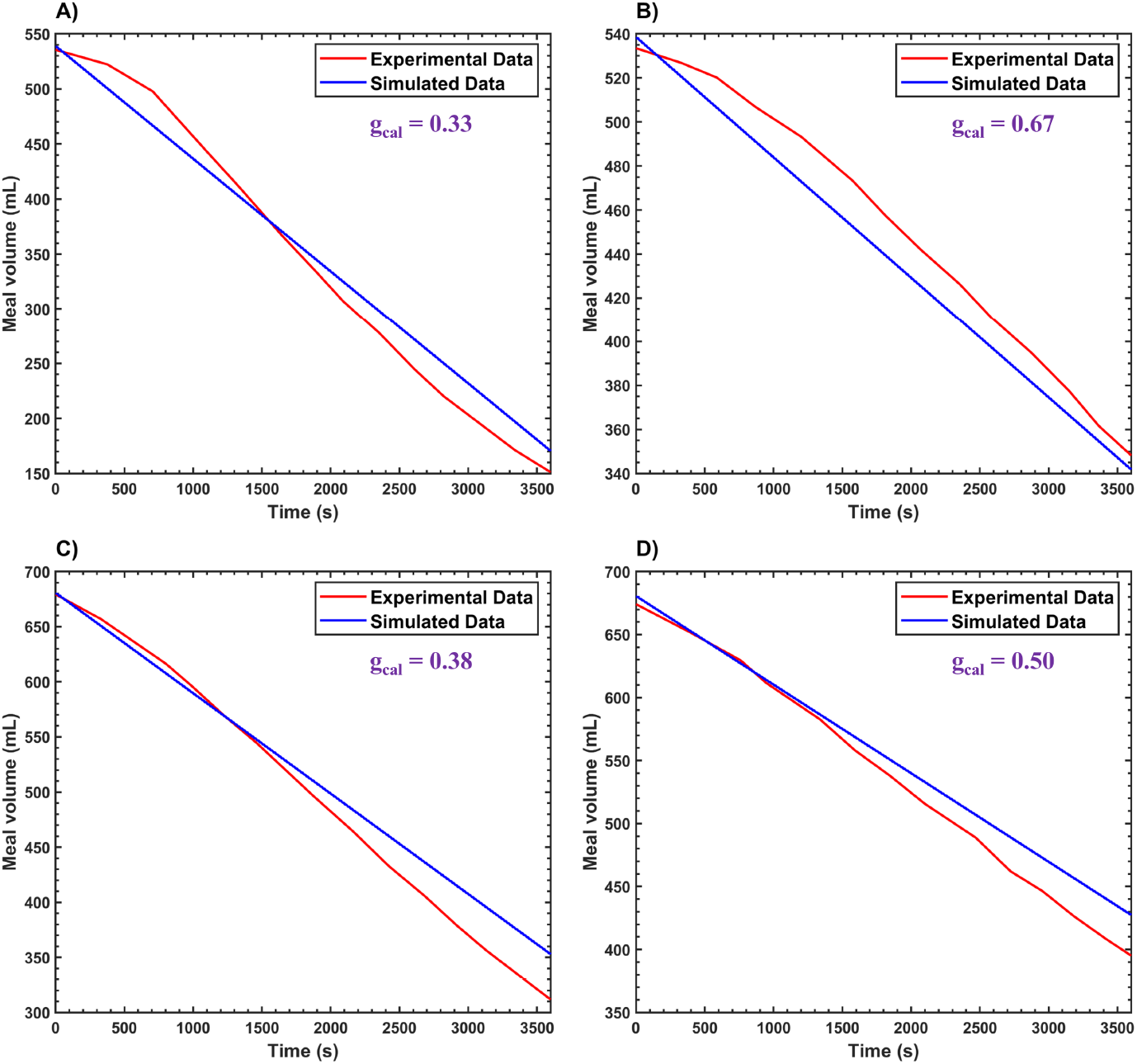
Gastric meal emptying plotted against time for gastric meals with different calorie contents for experimental [95] and simulated data, corresponding to *g*_*cal*_ values of 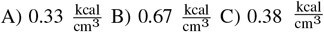 and 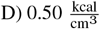

The emptying profiles show that gastric emptying rates are slower for meals with higher *g*_*cal*_ values compared to lower ones. This is because the <u>PS</u> modulates the rate of gastric emptying based on the caloric content of the meal, as discussed in Section <u>3.3.</u>

## 4. Conclusion

The model developed in this study simulates the vago-vagal loop that controls gastric function using a computationally efficient compartmental framework. This framework relies solely on <u>ODEs</u> and algebraic equations, as proposed in our prior work by Fernandes et al., 2024 [19]. The computational efficiency of the model enables it to be executed on a standard home laptop, making it highly accessible. Unlike previous efforts, which have modeled specific components of the gut-brain axis, this study represents the first attempt to comprehensively model the entire autonomic nervous system regulating gastric function. Previous mathematical models have addressed areas such as gastric cell electrophysiology [77, 115, 116], gastric whole-organ modeling [19, 24, 39, 117], enteric nerve physiology [118, 119, 120, 121, 122] and enteric neural inputs driving distal antrum phasic contractions [69]. This study presents a lumped yet detailed model of the vago-vagal loop, designed with potential applications in treating <u>GI</u> diseases through control theory. This is particularly significant given the prevalence of gut-brain axis dysfunction in conditions such as functional dyspepsia [123].

In addition to modeling the neural pathways, this study introduces advancements in the representation of gastric organ dynamics compared to our previously developed model [19]. Specifically, the new model accounts for dynamic changes in gastric volume rather than relying on a fixed geometry. However, the fluid mechanics within the stomach have been simplified to prioritize the effects of neural pathways on gastric motility. For instance, the antrum is now represented as a single compartment, unlike the three-compartment representation used in the prior model [19]. This simplification limits the detailed resolution of fluid dynamics. Future work aims to extend the compartmental framework to include more detailed fluid mechanics, potentially improving the physiological accuracy of the model.

In the future, several promising directions exist for the continuation of this research. Validation of specific model components against experimental data is an important next step. Additionally, the model could be adapted for use in vagal nerve stimulation therapy by incorporating it into a model-based closed-loop controller [124, 125, 126]. Such an application would leverage the computational efficiency of the model and detailed representation of neural-gastric interactions to provide novel therapeutic approaches for <u>GI</u> diseases.

## 5. Code Availability

The MATLAB code supporting this paper is available on GitHub at: https://github.com/shanferns/Gut-brain-axis-compartment-model.git

## 7. Acknowledgements

This work was supported by the National Institutes of Health, USA, Grant OT2OD030535, under the Stimulating Peripheral Activity to Reduce Conditions (SPARC) program; the John C. Chen graduate fellowship from Chemical and Biomolecular Engineering at Lehigh University for Mr. S. Fernandes; and a Faculty Innovation Grant (FIG) from Lehigh University. The authors would like to thank Dr. Oluwasanmi Adeodu for his valuable discussions and insightful suggestions, which greatly contributed to the mathematical modeling approach for representing the neural pathway of the gut-brain axis.

