## Supplemental Text for "A Compartmental Model for Simulating the Gut-Brain Axis in Gastric Function Regulation"

### 8. Supplementary Material

#### 8.1. Applying the MMEHC equation to model neuronal firing effects on smooth muscle responses

To apply the MMEHC equation, neurotransmitter release following neuronal firing is analyzed. Specifically, the firing dynamics of a DMV neuron projecting to the stomach is modeled. The action potential of the DMV neuron is simulated using Hodgkin–Huxley-type equations, with most ion channel gating kinetics following the framework outlined in Briant et al., 2014 [127]. Modifications were introduced to simplify the model reprenting of the DMV neuron.

The calcium concentration at the synaptic cleft, resulting from DMV neuron action potentials, is modeled using equations from Erler et al., 2004 [128]. The corresponding neurotransmitter release, driven by synaptic calcium concentration, is modeled using a mathematical framework from Briant et al., 2015 [53]. The relationship between DMV action potential firing frequency and normalized neurotransmitter release is then plotted, and the MMEHC equation is fitted to these data for analysis.

For modeling neurotransmitter release and its effect on smooth muscle response—such as changes in contractile force—the MMEHC equation can be applied based on Briant et al., 2015 [53]. The plots in this study demonstrate that these dynamics follow non-linear saturation curves, which the MMEHC equation effectively captures. Moreover, the relationship between neurotransmitter release and smooth muscle response resembles a lumped ligand-receptor binding mechanism, a type of interaction that the MMEHC equation is well-suited to model.

##### 8.1.1. Neuron modeling: Action potential in the DMV

The governing equation for the membrane voltage of the DMV neuron, $V_{m,DMV}$, is given by

$$\frac{dV_{m,DMV}}{dt} = \frac{-I_{ion,DMV} + I_{stim,DMV}}{C_{m,DMV}} \tag{S.1}$$

Here, $I_{ion,DMV}$ represents the total ionic current in the DMV neuron, computed as the sum of sodium, potassium, calcium, and leak currents, derived from the dynamics of voltage-gated channels. $I_{stim,DMV}$ denotes the external stimulation current applied to the DMV neuron. Most ionic channels are modeled following the approach of Briant et al., 2014 [127]. However, certain parameters, such as the conductance of specific ion channels, were manually adjusted to better fit the experimental action potential data of DMV neurons projecting to the stomach, as reported by Browning et al., 2005 [129].

$$I_{ion,DMV} = I_{Na,DMV} + I_{KDR,DMV} + I_{Pas,DMV} + I_{KCa,DMV} + I_{CaL} + I_{CaN} \tag{S.2}$$

The leak current, $I_{Pas,DMV}$, modeled similarly to Briant et al., 2014 [127], is given by

$$I_{Pas,DMV} = g_{Pas}(V_{m,DMV} - E_L) \tag{S.3}$$

The calcium activated potassium current, $I_{KCa,DMV}$, modeled similarly to Briant et al., 2014 [127], is denoted by

$$I_{KCa,DMV} = g_{KCa} o_{KCa} (V_{m,DMV} - E_K) \tag{S.4}$$

The delayed rectifier current, $I_{KDR,DMV}$, modeled similarly to Briant et al., 2014 [127], is represented by

$$I_{KDR,DMV} = g_{KDR} n_{KDR}^3 l_{KDR} (V_{m,DMV} - E_K) \tag{S.5}$$

The sodium current, $I_{Na}$, is modeled similarly to Briant et al., 2014 [127]. However, the channel representation was simplified to a traditional sodium channel with only two gating variables. The activation ($m_{Na,DMV}$) and inactivation ($h_{Na,DMV}$) gating variables follow the same formulation as described by Briant et al., 2014 [127]. The sodium current equation is given by

$$I_{Na} = g_{Na} m_{Na,DMV}^3 h_{Na,DMV} (V_{m,DMV} - E_{Na}) \tag{S.6}$$

The N-type calcium current, $I_{CaN}$, modeled similarly to Briant et al., 2014 [127], is denoted by

$$I_{CaN} = -g_{CaN} m_{CaN}^2 \left( \frac{0.001}{0.001 + [Ca^{2+}]_{i,DMV}} \right) 12.5 \left( 1 - \frac{[Ca^{2+}]_{i,DMV}}{[Ca^{2+}]_{o,DMV}} \right) \exp\left( \frac{V_{m,DMV}}{12.5} \right) \operatorname{erf}\left( \frac{V_{m,DMV}}{12.5} \right) \quad \text{(S.7)}$$

The L-type calcium current, $I_{CaL}$, is modeled similarly to Briant et al., 2014 [127]. However, the intracellular calcium concentration in the DMV neuron is scaled by a factor of 5 to adjust the calcium concentration to match its contribution to the total current in the neuron. The equation is given by

$$I_{CaL} = -g_{CaL} m_{CaL}^2 \left( \frac{0.001}{0.001 + [Ca^{2+}]_{i,DMV}} \right) 12.5 \left( 1 - \frac{5[Ca^{2+}]_{i,DMV}}{[Ca^{2+}]_{o,DMV}} \right) \exp\left( \frac{V_{m,DMV}}{12.5} \right) \operatorname{erf}\left( \frac{V_{m,DMV}}{12.5} \right) \quad \text{(S.8)}$$

The potassium activated calcium current, $I_{KCa,DMV}$, modeled similarly to Briant et al., 2014 [127], is denoted by

$$I_{KCa,DMV} = g_{KCa,DMV} o_{KCa} \left( V_{m,DMV} - E_K \right) \quad \text{(S.9)}$$

The intracellular calcium concentration in the DMV neuron, $[Ca^{2+}]_{i,DMV}$, is modeled using a simplified approach based on Seydewitz et al., 2017 [130]. However, instead of using a Gaussian equation, the conversion from calcium currents to intracellular concentration is derived from Schild et al., 1994 [131]. The equation is given by

$$\frac{[Ca^{2+}]_{i,DMV}}{dt} = -\frac{(I_{CaL} + I_{CaN}) A_{DMV}}{F Z_{Ca} V_{DMV}} - k_{\infty}([Ca^{2+}]_{i,DMV} - [Ca^{2+}]_{rest,DMV}) \quad \text{(S.10)}$$

where

$$V_{DMV} = \frac{4}{3} \pi r_{DMV}^3 \quad \text{(S.11)}$$

$$A_{DMV} = 4 \pi r_{DMV}^2 \quad \text{(S.12)}$$

Here, $r_{DMV}$ represents the cell radius, $V_{DMV}$ represents the cell volume, and $A_{DMV}$ denotes the cell surface area. The valence of calcium ions is given by $Z_{Ca}$, and $F$ is Faraday's constant. The resting intracellular calcium concentration in the DMV neuron is denoted as $[Ca^{2+}]_{rest,DMV}$.

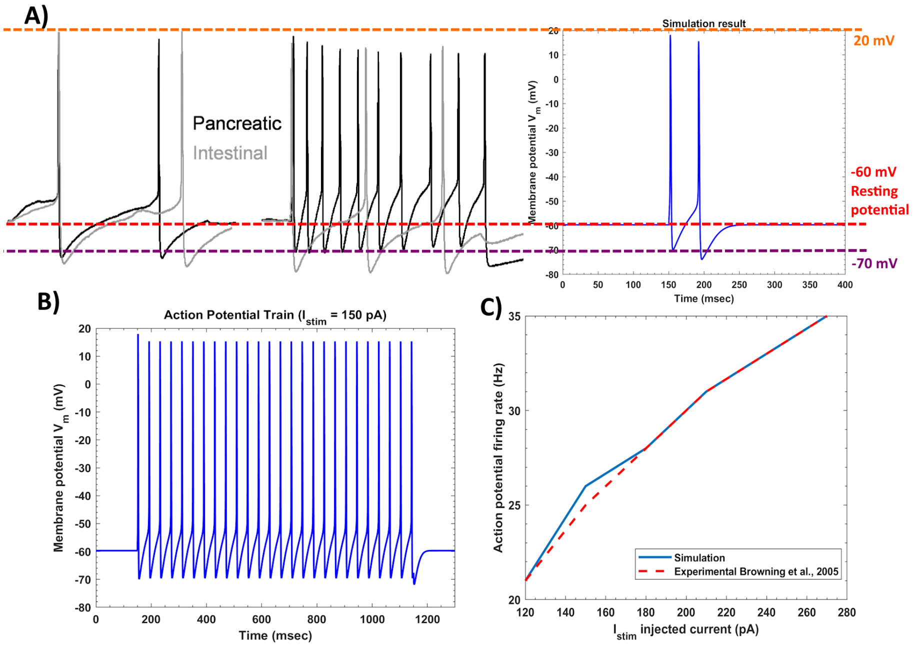


Figure 9: Action potential of DMV neurons projecting to the stomach A) Comparison of shape of action potentials between Experimental [129] and simulated by the models proposed in this section B) train of action potentials simulated for external stimulus current $I_{stim} = 150$ pA C) Comparison of experimental [129] and simulated action potentials trains for different $I_{stim}$ values

Table 3: Parameters for DMV neuron action potential

| Parameter | Value | Unit | Reference |
|---|---|---|---|
| $g_{Pas}$ | 0.26 | $\mu S\,cm^{-2}$ | Chosen |
| $g_{Na}$ | 40 | $\mu S\,cm^{-2}$ | Chosen |
| $g_{CaL}$ | 1 | $\mu S\,cm^{-2}$ | Chosen |
| $g_{CaN}$ | 8 | $\mu S\,cm^{-2}$ | Chosen |

*Continued on next page*

*Continued from previous page*

| Parameter | Value | Unit | Reference |
|---|---|---|---|
| $g_{KDR}$ | 2 | $\mu S\,cm^{-2}$ | Chosen |
| $g_{KCa}$ | 10 | $\mu S\,cm^{-2}$ | Chosen |
| $[Ca^{2+}]_{rest,DMV}$ | 50 | nM | [132] |
| $r_{DMV}$ | 12 | µm | [129] |
| $Z_{ca}$ | 2 | mV | [131] |
| $E_K$ | -80 | mV | Chosen |
| $E_L$ | -40 | mV | [127] |
| $E_{Na}$ | 20 | mV | Chosen |
| $k_\infty$ | 2 | $ms^{-1}$ | Chosen |
| $C_{m,DMV}$ | 1 | $\mu F\,cm^{-2}$ | [132] |

*8.1.2. Calcium concentration in the synaptic cleft*

The calcium concentration in the synaptic cleft, which is essential for neurotransmitter release from vesicles in response to an action potential in the DMV neuron, is modeled using a mathematical framework developed by Erler et al., 2004 [128]. The parameters governing this model can be found in their study.

The time-dependent single-channel open probability, $g_v$, is described by

$$\frac{dg_v}{dt} = \frac{\hat{g}_v - g_v}{\tau} \tag{S.13}$$

where $\tau$ is the time constant.

The calcium concentration in the synaptic cleft of the DMV neuron, denoted as $[Ca^{2+}]_{free}$, evolves according to

$$\frac{d[Ca^{2+}]_{free}}{dt} = \left(\frac{[G]}{Z_{Ca}F}\right)(J_i - J_e + L)\,\frac{1}{1 + T_{en} + T_{ex}} \tag{S.14}$$

The steady-state open probability, $\hat{g}_v$, follows a sigmoidal relationship

$$\hat{g}_v = \frac{1}{\exp\left(\frac{V_h - V_{m,DMV}}{\epsilon}\right) + 1} \tag{S.15}$$

where $\epsilon$ represents the steepness of activation, and $V_h$ is the half-activation voltage.

The inward current, $I_{\text{open}}$, associated with channel opening due to an action potential in the DMV neuron, is defined as

$$I_{\text{open}} = \begin{cases} 0, & V_{m,DMV} > \bar{V}_c \\ \bar{g}_v(\bar{V}_c - V_{m,DMV}), & \text{otherwise} \end{cases}$$

where $\bar{V}_c$ is given by

$$\bar{V}_c = \left(\frac{RT}{zF}\right) \ln\left(\frac{[Ca]_{ext}}{[Ca^{2+}]_{free}}\right) - \Delta V_{eff} \tag{S.16}$$

The calcium influx current density, $J_i$, is defined as

$$J_i = \rho_v g_v I_{\text{open}} \tag{S.17}$$

For the complete set of model equations, refer to Erler et al., 2004 [128]. The simulated synaptic cleft calcium concentration for different action potential frequencies in the DMV neuron is presented in Fig. 10.

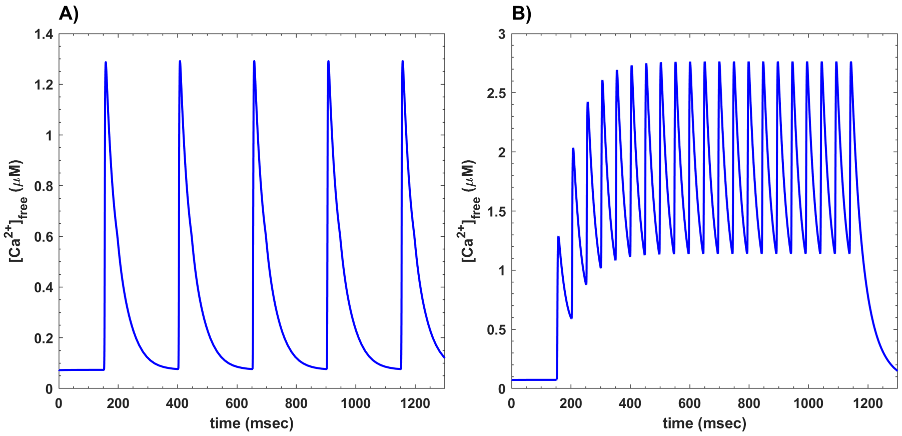


Figure 10: Calcium concentration in the DMV neuron synapse $[Ca^{2+}]_{free}$ for different firing frequency of DMV neuron action potential firing frequency A) 4 Hz B) 21 Hz

#### 8.1.3. Neurotransmitter release

The concentration of neurotransmitter release, $[NA]$, in response to calcium concentration at the synaptic cleft, $[Ca^{2+}]_{free}$, is simulated using the model from Briant et al. (2015) [53]. The parameters for this model are detailed in that study. However, the calcium ion binding rate constant, $k_b$, was varied with values of $k_b = 10^{14}\frac{1}{\text{mM}^4\text{ms}}$, $k_b = 10^{13}\frac{1}{\text{mM}^4\text{ms}}$, and $k_b = 10^{12}\frac{1}{\text{mM}^4\text{ms}}$. The corresponding simulation results are shown in Fig. 11. This variation was introduced to assess the sensitivity of neurotransmitter release to different rates of calcium-mediated protein binding.

The equations governing the Briant et al. (2015) [53] model are:

$$\frac{d[F_A]}{dt} = k_b(F_{max} - [F_A] - [V_A])[Ca^{2+}]_{free}^4 - k_u[F_A] - k_1[F_A]V + k_2[V_A] \quad \text{(S.18)}$$

$$\frac{d[V_A]}{dt} = k_1[F_A]V - (k_2 + k_3)[V_A] \quad \text{(S.19)}$$

$$\frac{d[N_A]}{dt} = Nk_3[V_A] - k_h[NA] \quad \text{(S.20)}$$

From Fig. 11, it can be observed that fitting a MMEHC curve to the simulated data—relating neuronal firing frequency to neurotransmitter release—demonstrates a close alignment with the expected behavior. This suggests that the MMEHC equation is an appropriate model for capturing this relationship.

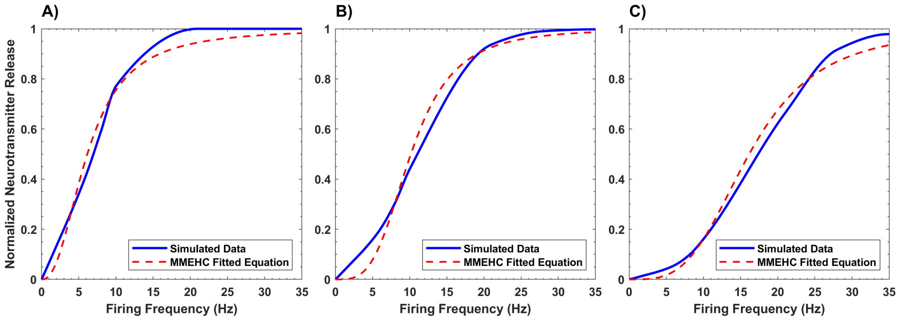


Figure 11: Comparison of simulated data from the model with MMEHC equation fit for normalized neurotransmitter release concentration as a function of DMV neuron firing frequency. (A) $k_b = 10^{14}\,\frac{1}{\mathrm{mM}^4\mathrm{ms}}$, (B) $k_b = 10^{13}\,\frac{1}{\mathrm{mM}^4\mathrm{ms}}$, (C) $k_b = 10^{12}\,\frac{1}{\mathrm{mM}^4\mathrm{ms}}$.

### *8.2. Efferent neuron and stomach compartment parameters*

#### *8.2.1. Fundus*

The MMEHC equation is used to model neurotransmitter release based on the firing frequency of inhibitory and excitatory neurons. Additionally, this equation modulates the relaxation or contraction response induced by neurotransmitter release in the fundus through signaling pathways. The relaxation or contraction response is computed as values ranging from 0 to 1, with the equations derived using the surface area of an open cylinder.

The relaxation response is given by

$$relax = \frac{r_{fin,1} - r_{min,1}}{r_{max,1} - r_{min,1}} \tag{S.21}$$

The contraction response is then computed as

$$contract = 1 - relax \tag{S.22}$$

Here, $r_{min,1}$ is determined by modeling the stomach as a cylinder and calculating its radius when the gastric volume is at a minimum (0.08 L). Similarly, $r_{max,1}$ is computed by assuming the stomach is cylindrical and calculating the radius at the maximum volume (1.2 L). The value of $r_{fin,1}$ is obtained using Eq. 15.

The parameters for the MMEHC equation were estimated using experimental data from [52, 56] to fit the response of cholinergic neuron firing frequency to neurotransmitter (Ach) release and the subsequent contraction response. Data from [59, 60, 61, 46] were used to fit the response of NANC neuron firing to neurotransmitter (NO and VIP) release and the corresponding muscle relaxation response.

For parameter fitting, MATLAB 'Curve Fitting' toolbox was utilized, employing the 'NonlinearLeastSquares' method with the 'Levenberg-Marquardt' algorithm. Manual adjustments were made to ensure the parameters were physiologically meaningful and produced the appropriate response.

The cholinergic and NANC pathway responses for the fundus are shown in Fig. 12. The response aligns well with experimental data reported in the literature [59, 133, 60, 52, 63]. The response for the cholinergic pathway successfully meets the expected contraction response at the desired firing frequency (Fig. 12 A)). For the NANC pathway, the relaxation response is initially dominated by NO when the inhibitory neuron firing frequency is below 0.5 Hz, after which the VIP pathway contributes to the relaxation response in the fundus. This behavior is consistent with previously reported findings [46, 48, 49, 50].

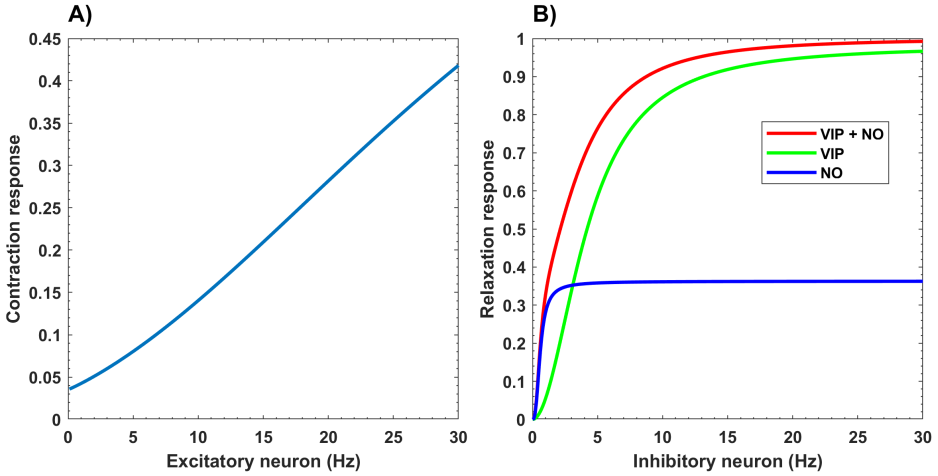


Figure 12: A) Cholinergic neuron firing frequency vs. fundus tissue contraction response when the tissue is fully relaxed ($f_{i,to,1}$ = 15 Hz) B) NANC neuron firing frequency vs. fundus tissue relaxation response. The blue line represents the response for the NO neurotransmitter signaling pathway, the green line represents the response for the VIP neurotransmitter signaling pathway, and the red line represents the combined effect of both pathways

In the fundus muscle tissue, the intracellular calcium concentration at resting state, $[Ca]^{2+}_{rest,1}$, is determined by setting the SMC membrane voltage to $-45$ mV [24] and using the Corrias and Buist, 2007 SMC model [77]. The $[MLCP]_{max,1}$ value, is assumed to be 7.5 μM, based on the value reported by Gajendiran and Buist, 2011 [28]. This value was chosen as it provides optimal responses for muscle relaxation mediated by NO and VIP neurotransmitter signaling.

The parameter $\beta_1$ for the fundus was adjusted to ensure that $r_{fin,1}$ values in Eq. 15 ranged between $r_{min,1}$ and $r_{max,1}$. The dimensionless constant $\alpha_1$ was computed to maintain $\lambda_{f,1}$ within the range of 1 to 1.7, which is the optimal limit reported by Panda and Buist, 2021 [31].

#### *8.2.2. Antrum*

To model efferent neuron firing in the antrum, data from Athavale et al., 2024 [69] was primarily used. This study [69] focused on the inhibitory and excitatory nuron firing responses of ICC and SMC via cholinergic, nitrergic, and purinergic pathways.

The data was analyzed based on factors such as ICC and SMC membrane voltage. Fractional values were obtained by normalizing ICC and SMC membrane voltage amplitudes against their baseline values. Similarly, the “slow wave” frequency was normalized against its baseline.

However, additional data were incorporated beyond Athavale et al., 2024 [69]. Specifically, to model excitatory pathway frequency, data from Forrest et al., 2006 [71] was used instead of Athavale et al., 2024, as the latter reported minimal changes in “slow wave” frequency at low cholinergic neuron firing rates. To model the impact of the NANC pathway on tissue stress modulation and contraction ratio, data from Kim et al., 2003 [72] was used. The contraction ratio $CR$ is computed as

$$CR = 1 - \frac{r_{fin,2}}{r_{ini,2}} \tag{S.23}$$

where $r_{fin,2}$ and $r_{ini,2}$ are obtained from Eq. 42. Additional data [74, 71, 73, 72, 62] were used to model neuron firing frequency, neurotransmitter release, and their effects on contraction intensity in the antrum.

The MMEHC equation parameters were fitted to model cholinergic and NANC responses in the antrum. The responses for excitatory and inhibitory neuron firing, their effects on contraction ratio, and ”slow wave” frequency are shown in Fig. 13. The results align with previous studies, where increasing excitatory neuron firing frequency leads to increased contraction ratio and ICC frequency, consistent with Athavale et al., 2024 [69]. Minimal increases in ICC frequency at low excitatory firing frequencies align with Forrest et al., 2006 [71]. Increasing inhibitory firing frequency reduces the contraction ratio while having little effect on ICC frequency, consistent with Kim et al., 2003 [72].

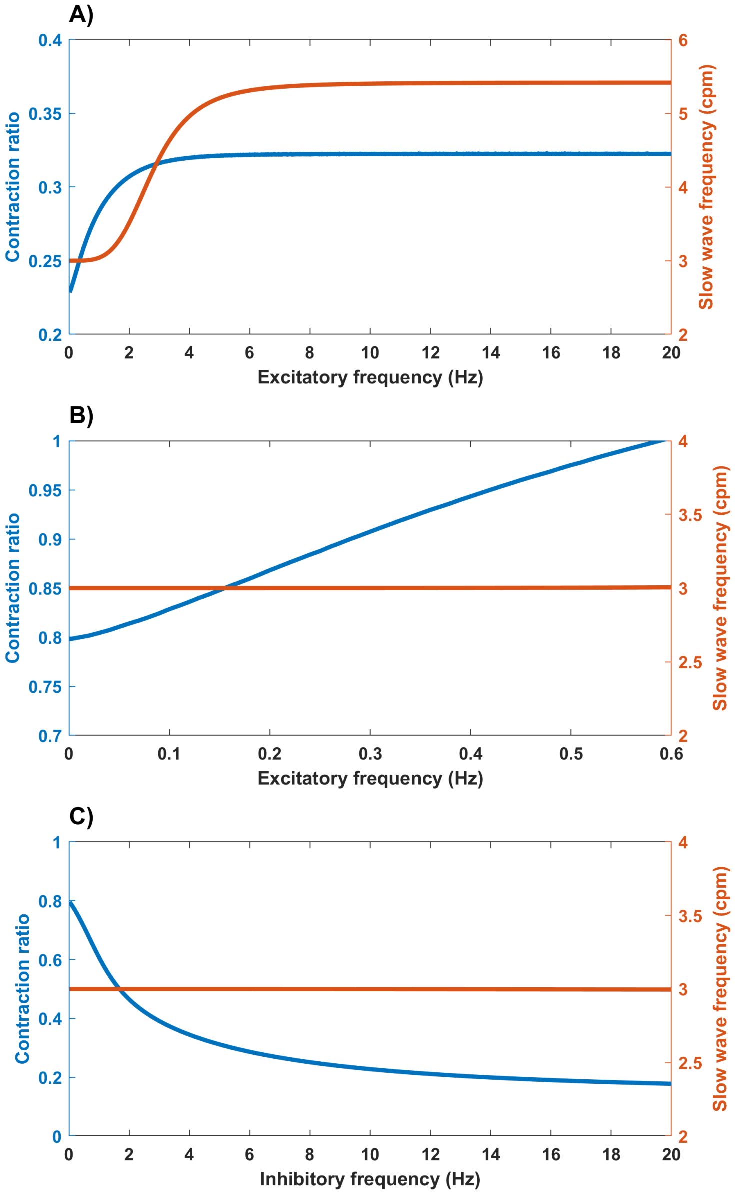


Figure 13: A) Excitatory neuron firing frequency vs. contraction ratio vs. ICC frequency when the tissue is relaxed ($f_{i,p,2}$ = 15 Hz). B) Excitatory neuron firing frequency vs. contraction ratio vs. ICC frequency. C) Inhibitory neuron firing frequency vs. contraction ratio vs. ICC frequency.

To compute tissue stretch, the hyperelastic component of the NLVM model was modified using a fifth-order polynomial equation, fitted to stress-stretch data from Panda and Buist, 2021 [31]. The choice of a polynomial approach follows the method of Panda and Buist, 2018 [82], with a fifth-order polynomial selected to capture the highly nonlinear stress-stretch response. The viscoelastic behavior remains modeled using parameters from our previous study [19].

Table 4: Coefficient values for principal stress $E_w$

| Coefficient | Value |
|---|---|
| $\mathcal{A}_4$ | $305.9792$ |
| $\mathcal{A}_3$ | $-2321.3779$ |
| $\mathcal{A}_2$ | $7068.6556$ |
| $\mathcal{A}_1$ | $-10773.4619$ |
| $\mathcal{A}_0$ | $8232.1226$ |

### *8.2.3. PS*

To model the tonic efferent response of the PS, a similar approach to that described in Section 8.2.1 was used. Due to the lack of data on the cholinergic response of the PS, data from cholinergic neurons in the fundus were used instead. This substitution was based on experimental findings indicating that the PS exhibits a contraction response to a given Ach concentration similar to that observed in the fundus [134, 135, 136].

To model the inhibitory response of the NANC pathway, data from Ishiguchi et al., 2000 [87] were used. However, the specific inhibitory neurotransmitter for the PS remains unclear [83]. As a result, NANC neuron stimulation was modeled as directly influencing inhibitory signaling without explicitly simulating neurotransmitter release.

Using these experimental data, parameters for the MMEHC equation were determined for both the cholinergic and NANC neuron pathways. The corresponding model response is presented in Fig. 14. The contraction and relaxation responses align well with experimental findings [134, 87].

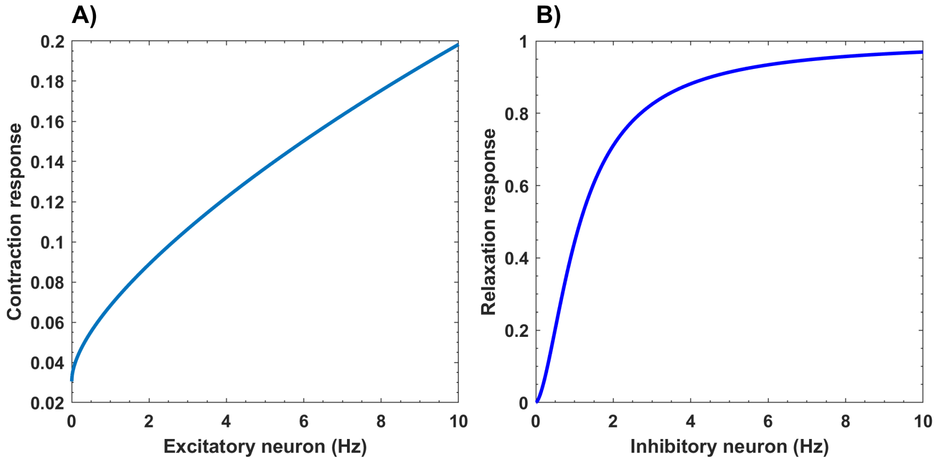


Figure 14: A) Cholinergic neuron firing frequency vs. PS tissue contraction response when the tissue is fully relaxed ($f_{i,to,3}$ = 10 Hz). B) NANC neuron firing frequency vs. PS tissue relaxation response.

To compute the resting intracellular calcium concentration in the PS ($[Ca]^{2+}_{rest,3}$), a method similar to that described in Section 8.2.1 was employed. However, the resting SMC membrane voltage in the PS is reported to be $-57$ mV [137, 138], and the corresponding resting intracellular calcium concentration was calculated accordingly.

The parameter $\beta_3$ for the PS was adjusted to ensure that $r_{fin,3}$ values in Eq. 15 remained within the range of $r_{min,3} = 0.075$ cm to $r_{max,3} = 0.48$ cm [138, 19, 139]. Additionally, the value of $\alpha_3$ was computed to maintain $\lambda_{f,3}$ within the range of 1 to 1.7, as discussed in Section 8.2.1.

To compute the gastric flow rate through the PS, $Q_{flow}$, a simplified equation is used. Here, $Q_{flow,max}$ represents the maximum flow rate, which occurs when the PS is fully open. The value of $Q_{flow,max}$ is derived from literature data for gastric emptying of a zero-calorie liquid with low viscosity, such as water [140, 141, 19]. For such a liquid, gastric emptying occurs at the fastest rate. This flow rate is then adjusted by a factor that accounts for the resistance to gastric flow, which depends on the sphincter radius and the degree of PS occlusion. To keep the model simple, a quadratic relationship is applied, consistent with literature on liquid flow through valves at different opening percentages [19, 142, 143, 144].

*8.3. Brainstem and intramural connections*

To model the correlation between afferent and efferent neuron signaling, Park et al., 2020 [97] established a relationship between their firing frequencies using a sigmoid function.

For the mechanoreceptor that influences antral contractions based on gastric volume, a similar approach was used as in Park et al., 2020 [97] to correlate afferent and efferent neuron firing. In this model, the mechanosensitive afferent neuron firing frequency, $f_{mech}$, was linked to the excitatory cholinergic efferent neuron firing frequency in the antrum, $f_{e,p,2}$. This correlation is justified because, at higher gastric volumes, stronger antral contractions are required to facilitate gastric emptying [109]. The antral contractions (terminal antrum) were maintained at approximately 78–80 % [19, 139]. Desired values for afferent-efferent neuron firing correlations were obtained to achieve these contraction levels and are plotted in Fig. 15 A). The sigmoid function from Park et al., 2020 [97] (Eq. 49) was then fitted to the data, as shown in Fig. 15 A).

For the chemoreceptor that influences PS occlusion based on gastric meal caloric content, a similar methodology was used. The intuition behind relating afferent and efferent firing frequencies, as suggested by Park et al., 2020 [97], was applied. Based on the desired gastric emptying flow rate, the afferent neuron firing frequency was used to regulate the efferent neuron firing rate, which, in turn, controlled the PS radius. The desired correlation data were obtained and plotted in Fig. 15 B). In this model, the chemosensitive afferent neuron firing rate, $f_{chem}$, was linked to the inhibitory efferent neuron firing rate, $f_{i,to,3}$, since the PS must relax in response to gastric nutrient content to regulate the emptying rate. Since the NANC pathway is responsible for PS relaxation, the pathway was incorporated accordingly. However, as seen in Fig. 15 B), the correlation plot exhibited strong nonlinearity, making the sigmoid function from Park et al., 2020 [97] unsuitable. Instead, a polynomial function (Eq. 50) was used to capture the afferent-efferent neuron firing relationship while following the intuitive approach from Park et al., 2020 [97].

For the intramural connection that regulates fundus distension to maintain IGP based on gastric meal volume (as discussed in Section 3.4), gastric volume was estimated using the volume equation for a cylinder with a fixed height [19, 145, 146], while varying the radius $r_{fin,1}$. The value of $r_{fin,1}$ was obtained from Eq. 15 by varying the NANC pathway firing frequency $f_{i,to,1}$, which is responsible for fundus relaxation. The resulting data, representing the desired correlation, was plotted in Fig. 15 C). To model this relationship, a piecewise polynomial function (Eq. 51) was fitted to determine the extent of fundus distension required for a desired gastric volume. The fitted piecewise polynomial function and the desired correlation are illustrated in Fig. 15 C).

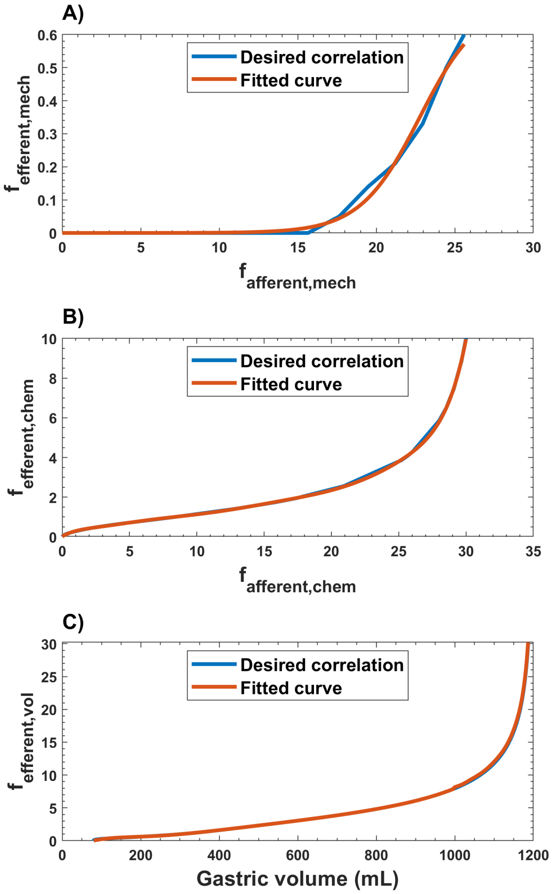


Figure 15: Desired correlation vs fitted curves for A) $f_{afferent,mech}$ (Hz) vs $f_{efferent,mech}$ (Hz) B) $f_{afferent,chem}$ (Hz) vs $f_{efferent,chem}$ (Hz) C) Gastric volume (mL) vs $f_{efferent,vol}$ (Hz)

Table 5: Parameter values for the mechanoreceptor brainstem interconnection

| Parameter | Value |
|---|---|
| $f_{min,mech}$ | 0 |
| $f_{max,mech}$ | 0.7 |
| $f_{mid,mech}$ | 22.77 |
| $k_{inter,mech}$ | 1.903 |

Table 6: Values of $k_{inter,chem,s-1}$

| $s$ | $k_{inter,chem,s-1}$ |
|---|---|
| 10 | $2.6286 \times 10^{-10}$ |
| 9 | $-3.2650 \times 10^{-8}$ |
| 8 | $1.7001 \times 10^{-6}$ |
| 7 | $-4.8147 \times 10^{-5}$ |
| 6 | $8.0636 \times 10^{-4}$ |
| 5 | $-8.1509 \times 10^{-3}$ |
| 4 | $4.9024 \times 10^{-2}$ |
| 3 | $-1.6874 \times 10^{-1}$ |
| 2 | $3.9815 \times 10^{-1}$ |
| 1 | $2.1430 \times 10^{-2}$ |

Table 7: Values of $k_{vol,h,s-1}$ and $k_{vol,l,s-1}$

| $s$ | $k_{vol,h,s-1}$ | $k_{vol,l,s-1}$ |
|---|---|---|
| 10 | $1.9882 \times 10^{-17}$ | - |
| 9 | $-1.9424 \times 10^{-13}$ | $-5.2753 \times 10^{-22}$ |
| 8 | $8.4307 \times 10^{-10}$ | $2.7237 \times 10^{-18}$ |
| 7 | $-2.1337 \times 10^{-6}$ | $-5.8880 \times 10^{-15}$ |
| 6 | $3.4702 \times 10^{-3}$ | $6.9430 \times 10^{-12}$ |
| 5 | $-3.7612$ | $-4.8382 \times 10^{-9}$ |
| 4 | $2.7166 \times 10^{3}$ | $2.0069 \times 10^{-6}$ |
| 3 | $-1.2609 \times 10^{6}$ | $-4.6750 \times 10^{-4}$ |
| 2 | $3.4126 \times 10^{8}$ | $5.8051 \times 10^{-2}$ |
| 1 | $-4.1033 \times 10^{10}$ | $-2.4861$ |

Here $V_{tot,o} = 997$
